# A Sterol-Binding Cavity Underlies Sterol Recognition and Differential Activation of the ABCG5/G8

**DOI:** 10.64898/2026.08.10.744064

**Authors:** Fatemeh Rezaei, Isra F. Omar, Danny Farhat, Zi-Wen Weng, Kadambari Vijay Sai, Qing-Fang Xiao, Yuan-Chih Chang, Shang-Te Danny Hsu, Jyh-Yeuan Lee

**Affiliations:** Department of Biochemistry, Microbiology and Immunology, Faculty of Medicine, University of Ottawa, 451 Smyth Rd, Ottawa, Ontario K1H 8M5, Canada; Translational and Molecular Medicine Program, Faculty of Medicine, University of Ottawa, 451 Smyth Rd, Ottawa, Ontario K1H 8M5, Canada; Institute of Biological Chemistry, Academia Sinica, 128 Academia Rd, Sec 2, Nankang, Taipei 11529, Taiwan; Institute of Biochemical Sciences, National Taiwan University, 1 Roosevelt Rd, Sec. 4, Daan, Taipei 10617, Taiwan; Biomedical Sciences Program, Faculty of Science, University of Ottawa, 30 Marie Curie Pvt, Ottawa, Ontario K1N 6N5, Canada; International Institute for Sustainability with Knotted Chiral Meta Matter (WPI-SKCM2), Hiroshima University, 1-3-1 Kagamiyama, Higashi-Hiroshima, Hiroshima 739-8531, Japan

**Keywords:** ABCG5/G8, ergosterol, cryo-electron microscopy, ATPase, molecular dynamic simulation

## Abstract

Sterol homeostasis depends on the coordinated regulation of endogenous cholesterol synthesis, dietary sterol absorption, and sterol excretion. The heterodimeric ATP-binding cassette sterol transporter ABCG5/G8 plays an important role in eliminating excess sterols by participating in reverse cholesterol transport and transintestinal cholesterol efflux. The molecular mechanism of sterol recognition and transport by ABCG5/G8 remains poorly understood. Here, we determined the cryo-electron microscopy (cryo-EM) structure of human ABCG5/G8 in complex with ergosterol. The structure revealed a sterol-binding site at the transmembrane domain (TMD) interface between the subunits ABCG5 and ABCG8, adjacent to the conserved aromatic clamp motif. Tyrosine 432 (Y432) on ABCG5, a key residue within the aromatic clamp, lies near the tetracyclic ring of ergosterol. Additionally, to assess the effect of different sterols on transporter activity, we performed molecular dynamic simulations and *in vitro* ATPase assays in the presence of cholesterol, cholesteryl hemisuccinate (CHS), and ergosterol. Ergosterol exhibited more favorable interactions with ABCG5/G8 and stimulated ATPase activity more effectively than either cholesterol or CHS, representing the first biochemical characterization of ABCG5/G8 activity in response to a non-cholesterol sterol. Furthermore, substitution of Y432 with the canonical phenylalanine in ABCG family abolished the differential ATPase response to ergosterol, with the mutant displaying similar activity levels in the presence of ergosterol and cholesterol. Together, our structural and biochemical findings reveal a conserved sterol-binding site within ABCG5/G8 and demonstrate direct evidence that distinct sterols differentially modulate ABCG sterol transporter activity and that the degenerative aromatic clamp motif in ABCG5 contributes to sterol-dependent functional selectivity.

## Introduction

Sterols are a diverse class of lipophilic compounds that constitute essential components of eukaryotic cell membranes (1). Due to their widespread occurrence in nature, sterols are present in the human diet as a diverse mixture that includes cholesterol, plant sterols, and ergosterol, derived from animal-, plant-, and fungal-based foods, oils, and insects (2–6). Among these, cholesterol is essential for human physiology, serving as a structural component of cellular membranes and as a precursor for steroid hormones and bile acids required for nutrient absorption, metabolism, and gene regulation (7). In addition to dietary intake, cholesterol is synthesized endogenously to maintain lipid homeostasis and ensure adequate supply under conditions of limited dietary intake (8). In humans, dietary cholesterol is absorbed relatively efficiently, whereas other sterols (xenosterols) are absorbed at much lower efficiencies, resulting in their markedly lower systemic abundance. This selective absorption represents an early regulatory checkpoint that shapes circulating sterol composition and contributes to whole-body sterol homeostasis (9,10).

At the systemic level, sterol homeostasis is maintained through coordinated regulation across multiple organs, primarily the intestine, liver, and peripheral tissues. Disruption of this balance, through either excessive intake or impaired clearance, can lead to pathological sterol accumulation and associated disease states (7,11). A central component of this regulatory network is the heterodimeric ATP-binding cassette G5/G8 (ABCG5/G8) transporter, which limits the accumulation of excess cholesterol and non-cholesterol sterols by mediating sterol efflux into the bile and the intestinal lumen via hepatobiliary secretion and transintestinal cholesterol efflux (TICE) pathways (12–14). Disruption of ABCG5/G8 function leads to sitosterolemia, an autosomal recessive disorder characterized by the accumulation of plant sterols, including sitosterol, campesterol, and stigmasterol, in plasma and tissues. This condition is also associated with progressive cholesterol accumulation and an increased risk of atherosclerosis (14–16).

Despite the well-established physiological role of ABCG5/G8 in sterol transport, the molecular basis of its interactions with diverse sterol substrates remains poorly understood, particularly with respect to non-cholesterol sterols. In this study, we determined the cryo-electron microscopy (cryo-EM) structure of ABCG5/G8 in a sterol-bound state. Structural analysis and molecular dynamics (MD) simulations identified an ergosterol-binding site adjacent to a conserved aromatic-clamp motif (previously named phenylalanine-highway motif) at the interface between the transmembrane domains (TMDs) of ABCG5/G8 (17). In parallel, we measured the sterol-coupled ATPase activity of ABCG5/G8 in the presence of cholesterol, ergosterol, or cholesteryl hemisuccinate (CHS), demonstrating that distinct sterol species differentially modulate transporter activity. Finally, using site-directed mutagenesis, we showed ABCG5 tyrosine 432 (ABCG5_Y432_), a key component of the aromatic clamp motif, as an important determinant of sterol-dependent regulation of ABCG5/G8 activity.

## Results

### Cryo-EM and MD analyses identified an ergosterol molecule bound between the TMDs of ABCG5/G8

In this study, using cryo-EM and single-particle analysis, we determined the structure of detergent-purified ABCG5/G8 in complex with ergosterol. To obtain a high-resolution structure, we combined two cryo-EM datasets comprising a total of 21,287 micrographs and 492,931 selected particles. The final density map achieved an overall resolution of 3.56 Å, according to the gold-standard Fourier shell correlation (GSFSC) 0.143 criterion (Fig. S2C). The 3D reconstruction displayed a high-quality density sufficient for model building of the TMDs at ∼3 Å resolution (Fig. S3), while the nucleotide-binding domains (NBDs) of both half-transporters were likewise well resolved (Fig. 1A and Fig. S2D).

**Figure 1.**
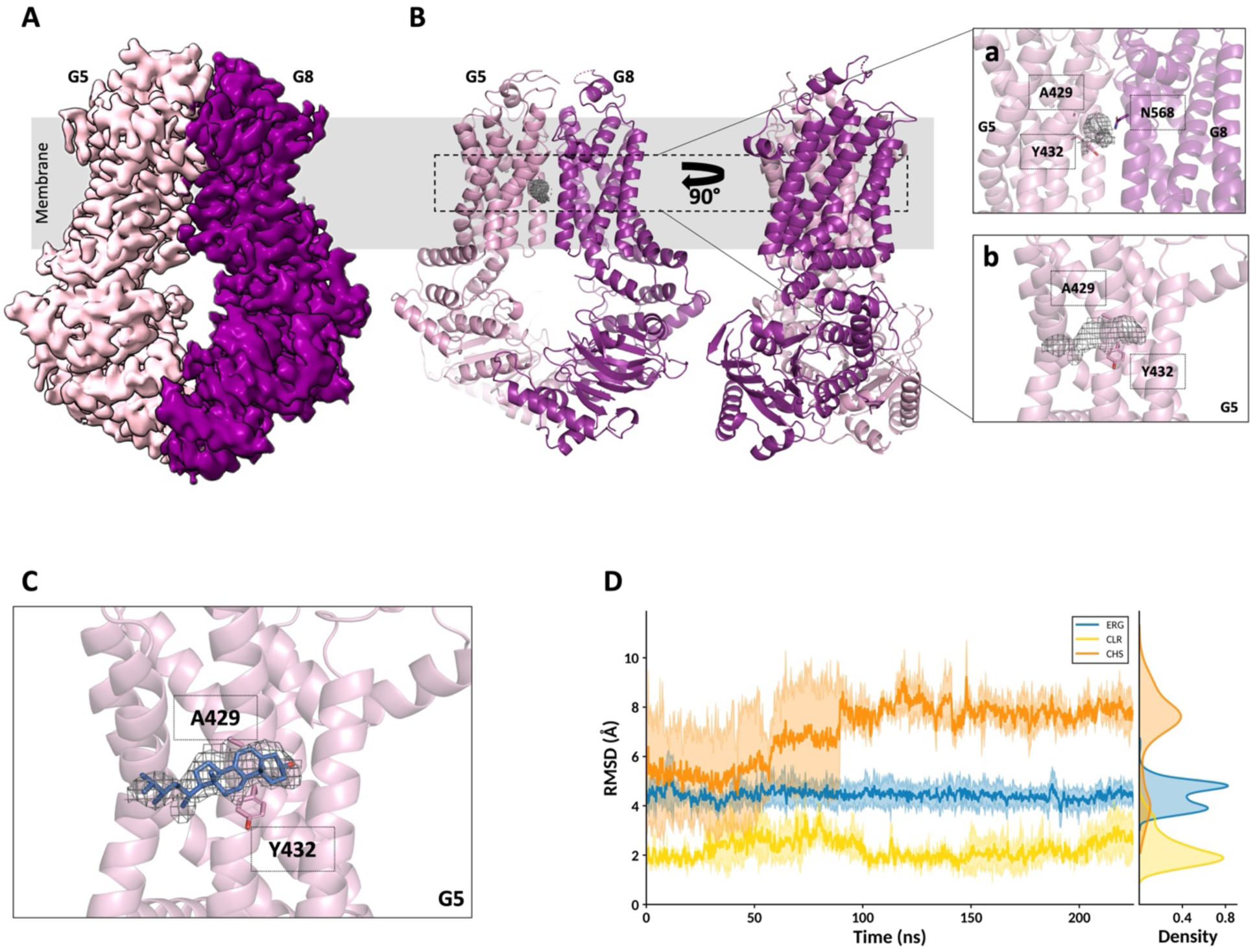
Cryo-EM structure of ABCG5/G8 showing a potential sterol-binding site buried between the TMDs of ABCG5 and ABCG8. **A and B.** Cryo-EM map and structure of ABCG5/G8 in the nucleotide-free (ATP-apo) and sterol-bound states. (**a and b)** The zoomed-in pictures of sterol-like density observed in between the TMDs of ABCG5/G8, adjacent to ABCG5_Y432_ and Alanine 429 (ABCG5_A429_). **C.** Ergosterol (blue) modeled into the observed cryo-EM density. **D.** RMSD profiles of ergosterol (blue), cholesterol (yellow), and CHS (orange) within the ABCG5/G8 binding site during 200 ns MD simulations across independent replicates.

Using a previously determined crystal structure of ABCG5/G8 (PDB ID: 8CUB) (17) as a starting model, we fitted and refined the model to better match the experimental electron density (Fig. 1B). Overall, the cryo-EM structure showed a root-mean-square deviation (RMSD) of 0.702 Å (1083 atoms) compared with the crystal structure. Similar to the crystal structure, ABCG5/G8 adopts an inward-facing conformation, with the TMDs open toward the cytoplasmic side, and retains an overall conserved architecture and key structural features (Fig. 1B). The statistics of cryo-EM data acquisition, processing, and refinement are summarized in Table S1, and the corresponding cryo-EM sample preparation and data processing workflows are shown in Fig. S1 and S2.

Following refinement of the model in the cryo-EM map, we identified a distinct and well-defined electron density located in a cavity at the interface between the TMDs of ABCG5 and ABCG8 (Fig. 1B, a and b). This density closely resembled the structure of a sterol molecule showing the sterol-nucleus and a tail feature and was oriented perpendicular to the membrane plane. Although the shape and size of this density were compatible with a cholesterol molecule, it was unlikely to represent cholesterol, given that cholesterol was not used at any stage of sample preparation. We therefore evaluated alternative sterol-like molecules that could potentially occupy this site, including ergosterol, the endogenous sterol produced by the Pichia pastoris system used for protein expression, and CHS, which was present during sample preparation.

For comparative analysis, ergosterol, CHS, and cholesterol were individually modeled and refined into the density (Fig. 1C and Fig. S4A, a and b). Ligand-map cross-correlation coefficients (CC) of 0.80, 0.55, and 0.75 were obtained for ergosterol, CHS, and cholesterol, respectively. Notably, the bulky succinate moiety of CHS was poorly accommodated within the observed density.

To further characterize sterol interactions within the binding site, we performed all-atom MD simulations to evaluate the interactions and stability of cholesterol, ergosterol, and CHS within this cavity of ABCG5/G8 over 200 ns for all conditions. MD simulations were performed using the most recent crystal structure (PDB ID: 8CUB) (17), with the missing chains completed using MODELLER. The protein was simulated in a membrane consisting of cholesterol and palmitoyl oleoyl phosphatidyl choline (POPC) at a ratio of 5:1 (mol:mol).

Consistent with the experimental data, MD simulations showed that cholesterol and ergosterol remained relatively stable in their docked poses, fluctuating between 2–5 Å RMSD across replicates, whereas CHS exhibited substantially greater positional mobility, with RMSD values reaching approximately 8 Å (Fig. 1D). The CHS deviations reflected a distinct translation away from the protein binding site where the density was observed and toward the cavity exit (Fig. S4B). Together, these observations suggest that CHS is unlikely to represent the bound ligand in this structure, and our results are more consistent with ergosterol occupying this site.

### Structural and computational analyses revealed molecular determinants of sterol recognition within the ABCG5/G8 transmembrane cavity

The observed sterol density in the cryo-EM map supports a defined binding pose of ergosterol within the binding cavity. Multiple orientations were evaluated by fitting ergosterol into the cryo-EM density; however, alternative poses resulted in lower ligand-map CC values and poorer agreement with the experimental density (Fig. S4C, a and b), supporting the assigned pose shown in Figure 1C. The sterol-binding pocket exhibits an amphipathic character, consistent with the mixed polar and hydrophobic properties of sterols and may contribute to the observed ligand orientation within the cavity (Fig. 2A). In this orientation, ergosterol is positioned with its 3β-hydroxyl group facing the dimer interface, where it lies within hydrogen-bonding distance of ABCG5 serine 538 (ABCG5_S538_) (Fig. 2B). In contrast, the sterol tetracyclic core and hydrocarbon tail extend toward the surrounding membrane environment, with ergosterol engaging ABCG5 through its α (smooth) face and ABCG8 through its β (rough) face (Fig. S4D).

**Figure 2.**
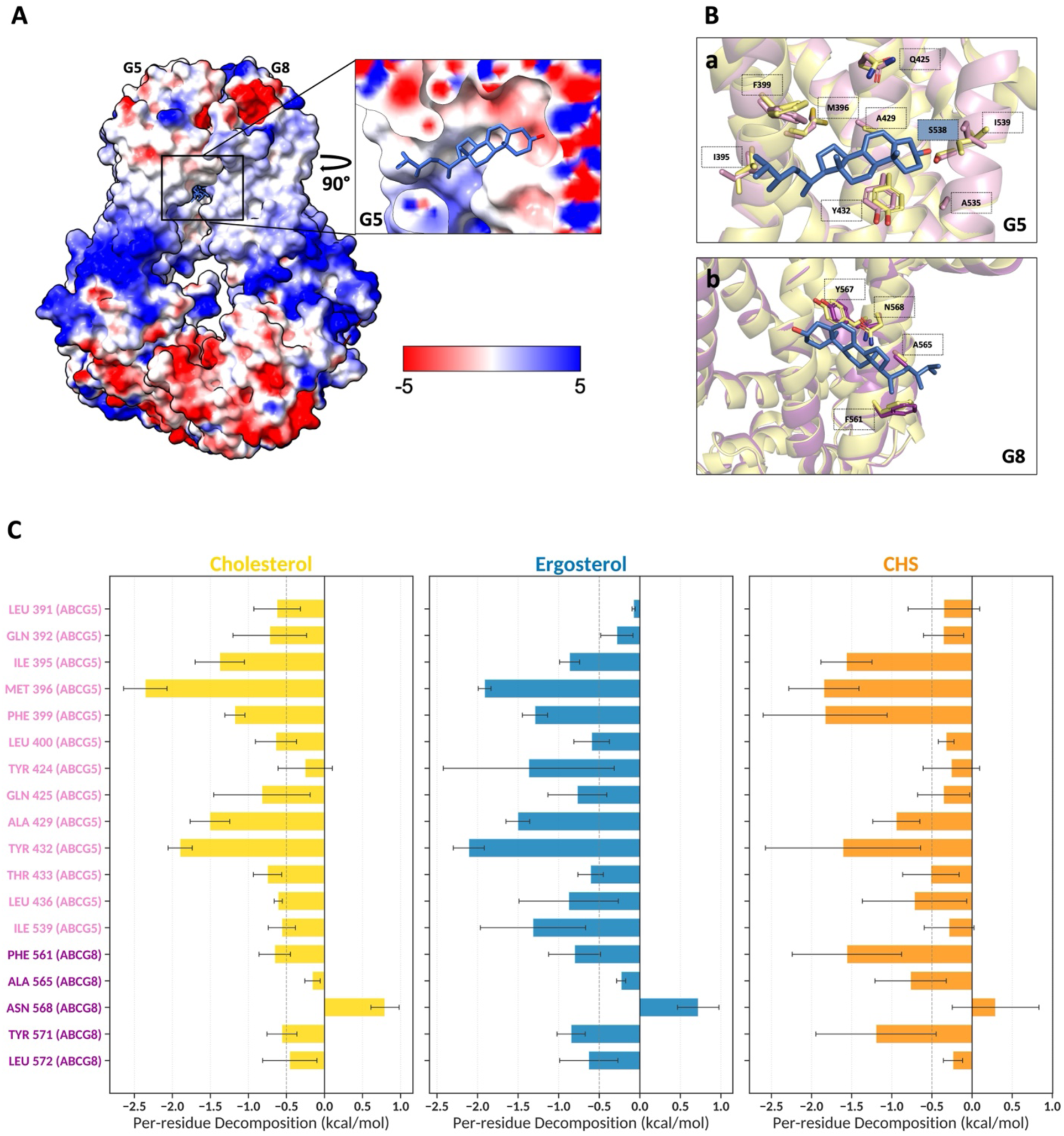
Structural features of the ABCG5/G8 sterol-binding pocket and molecular determinants of sterol recognition. **A.** Electrostatic surface map of ABCG5/G8. The zoomed-in view highlights the electrostatic potential within the binding pocket surrounding the observed bound ligand (ergosterol shown in blue). **B.** Alignment of the cryo-EM structure of ABCG5/G8 bound to ergosterol (ergosterol shown in blue; ABCG5 in pink and ABCG8 in purple) with the crystal structure (PDB ID: 8CUB; shown in pale yellow), which is apo at this binding site. **(a)** The alignment highlights conformational changes in residues forming the potential sterol-binding pocket in ABCG5. Serin 538 (highlighted in the blue box) is positioned 3.5 Å from the hydroxyl group of ergosterol. **(b)** The same structural alignment focusing on ABCG8, showing residue movements within the corresponding binding pocket. **C.** Per-residue binding free energy decomposition from the MD simulations of ABCG5/G8 in the presence of ergosterol (blue), cholesterol (yellow), or CHS (orange) over 200 ns MD simulations across independent replicates.

Alignment of the cryo-EM structure with our previously determined crystal structure, which is apo at this binding site (17), revealed that sterol binding induces conformational rearrangements within the binding pocket (Fig. 2B, a and b). These changes consisted primarily of side-chain movements and rotamer rearrangements among residues lining the cavity, with the most pronounced rotamer changes occurring in ABCG5 residues that form the binding pocket, together with ABCG8 asparagine 568 (ABCG8_N568_).

Per-residue binding free energy decomposition from the MD simulations of ABCG5/G8 in the presence of ergosterol, cholesterol, or CHS revealed substantial energetic contributions from TMH1 and TMH2 of ABCG5, as well as TMH2 of ABCG8, to sterol stabilization within the binding pocket. Among the residues analyzed, ABCG5_Y432_ and methionine 396 (ABCG5_M396_) consistently exhibited the most favorable binding free energy contributions for all three sterols, whereas ABCG8_N568_ showed the least favorable contribution to ligand binding (Fig. 2C). ABCG5_Y432_ and ABCG8_N568_ were previously identified as co-evolving residues, suggesting a conserved functional relationship between these positions (18).

MD simulation analysis further revealed sterol-dependent energetic contributions for several residues within the binding pocket (Fig. 2C). Overall, the residues exhibited greater variability in their energetic contributions during CHS-bound simulations than during cholesterol- or ergosterol-bound simulations. An exception was observed for ABCG5 tyrosine 424 (ABCG5_Y424_), which displayed substantially more favorable binding free energy contributions and greater energetic fluctuations in the presence of ergosterol than in the presence of cholesterol or CHS. ABCG5_Y424 and Y432_ are key components of a conserved aromatic clamp motif on ABCG5 (Fig. S5A)(17).

Distinct interaction patterns were also observed for ABCG5 isoleucine 395 (ABCG5_I395_) and isoleucine 539 (ABCG5_I539_). Relative to cholesterol- and CHS-bound simulations, ergosterol exhibited less favorable binding free energy contributions at ABCG5_I395_ but more favorable contributions at ABCG5_I539_ (Fig. 2C). Because ABCG5_I395 and I539_ define the entrance and distal end of the binding cavity, respectively, this shift in energetic contributions indicates that ergosterol is less stabilized near the cavity entrance and more favorably accommodated deeper within the pocket. Interestingly, these residues also showed the most pronounced side-chain rearrangements upon sterol binding relative to the previously determined crystal structure (Fig. 2B) (17).

### Assessment of ABCG5/G8 ATPase Activity Reveals Differential Modulation by Distinct Sterols

Although ABCG5/G8 is known to interact with multiple sterol substrates, previous *in vitro* measurements of its activity have been largely restricted to cholesterol and sterol mimetics, like CHS and bile acids (19–22). In this study, we optimized our previously established colorimetric citrate bismuth-based ATPase assay to enable activity measurements of wild-type ABCG5/G8 (ABCG5_WT_/G8_WT_) in the presence of non-cholesterol sterols, such as ergosterol (19). Using this approach, we directly measured ABCG5/G8-mediated ATP hydrolysis in the presence of cholesterol, CHS, and ergosterol. Under identical assay conditions containing 1.5% sodium cholate and 2 mg/mL soy phosphatidylcholine (soy PC), all three sterols stimulated ABCG5/G8 ATPase activity; however, ergosterol induced a stronger stimulation of ATP hydrolysis compared with cholesterol and CHS.

Under saturated ATP conditions (5 mM) and 0.25 mM sterols, time-course ATPase assays (0–20 minutes) showed that ATP hydrolysis by ABCG5/G8 was linear over the first 8 minutes. Initial rates were therefore calculated from the 2–8 minute interval (Fig. 3A). In the absence of sterol or CHS, the basal ATPase activity of ABCG5/G8 was 216 ± 72 nmol·min⁻¹·mg⁻¹ (mean ± SD from three independent ATPase assays; n=3), similar to previously reported values (19). This activity was increased by more than 2-fold (n=5) and 1.5-fold (n=4) in the presence of cholesterol and CHS, respectively. However, there was no significant difference between the rates achieved with cholesterol and CHS, indicating comparable levels of stimulation at a sterol concentration of 0.25 mM. On the other hand, in the presence of ergosterol, the ATPase activity was increased by more than 3.5-fold (n=7). This not only represents a rate enhancement of more than a 1.5 relative to CHS- or cholesterol-stimulated conditions, but it also represents the highest ABCG5/G8 ATPase activity reported to date (19–21). As a control, the catalytically impaired mutant ABCG5_WT_/G8_G216D_ was tested in the presence of ergosterol and exhibited substantially reduced ATPase activity compared with ABCG5_WT_/G8_WT_ under all sterol-stimulated conditions (Fig. 3A).

**Figure 3.**
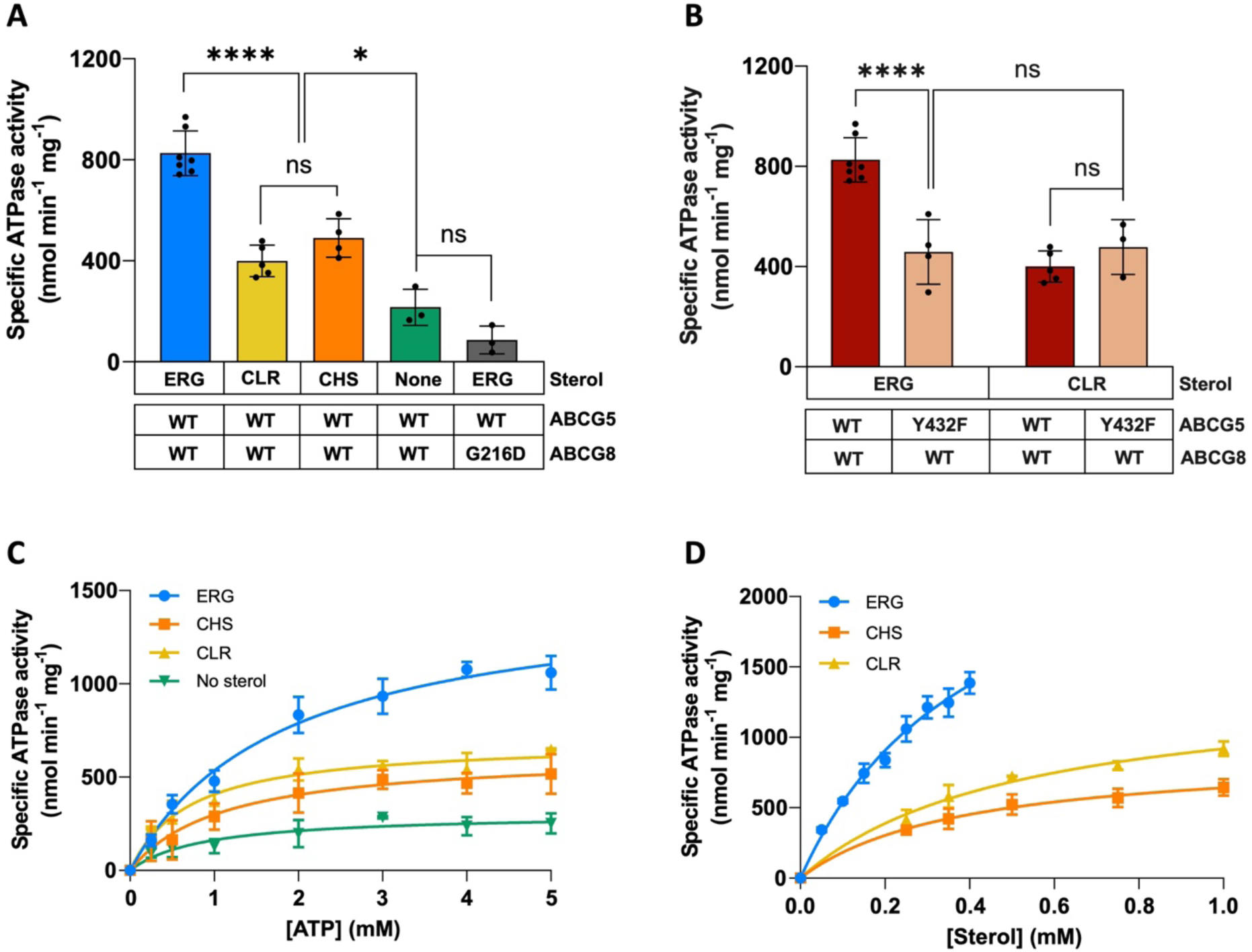
ATP- and sterol-dependent modulation of ABCG5/G8 ATPase activity. **A.** ATPase activity of ABCG5_WT_/G8_WT_ measured over the 2–8 min interval in the presence of ergosterol, cholesterol, or CHS, compared with the sterol-free condition and the catalytically inactive ABCG5_WT_/G8_G216D_ mutant. **B.** ATPase activity of ABCG5_Y432F_/G8_WT_ measured over the 2–8 min interval in the presence of ergosterol or cholesterol, compared with the corresponding ergosterol- and cholesterol-stimulated ATPase activities of ABCG5_WT_/G8_WT_ shown in panel A. Statistical significance in panels A and B was determined using one-way ANOVA with multiple comparisons. ns, not significant; *P < 0.05; ****P < 0.0001. **C.** ATP concentration-dependent ATPase activity of ABCG5_WT_/G8_WT_ measured in the presence of 0.25 mM ergosterol, cholesterol, CHS, or under sterol-free conditions. **D.** Sterol concentration-dependent ATPase activity of ABCG5_WT_/G8_WT_ measured in the presence of 5 mM ATP and increasing concentrations of ergosterol, cholesterol, or CHS. Ergosterol was insoluble above 0.4 mM under the current assay conditions.

To determine whether residues within the ergosterol-binding cavity contribute to sterol-dependent activation of ABCG5/G8, we focused on ABCG5_Y432_ (an important residue involved with the substrate density in the cryo-EM map) by replacing tyrosine with the conserved aromatic phenylalanine residue (ABCG5_Y432F_) found at the equivalent position in other ABCG transporters (17) (Fig. S5A). The ATPase activity of the ABCG5_Y432F_/G8_WT_ mutant was then assessed in the presence of ergosterol or cholesterol. For each sterol condition, mutant activity was compared with ABCG5_WT_/G8_WT_ under the corresponding condition (Fig. 3B). In the presence of ergosterol, the mutation resulted in a significant 1.8-fold reduction in ATPase activity (n=4), whereas in the presence of cholesterol, the mutation resulted in a non-significant change (n=3). Importantly, the mean rates achieved by the ABCG5_Y432F_/G8_WT_ mutant in either sterol-coupled conditions were almost identical to one another. ABCG5_Y432_ seems critical for ergosterol-binding-dependent activation of the transporter.

### Kinetic analysis reveals sterol-dependent modulation of ABCG5/G8 ATPase activity

To further characterize how different sterols modulate ATP hydrolysis, ATP concentration-dependent kinetic assays were performed in the presence of 0.25 mM ergosterol, cholesterol, or CHS, with ATP concentrations ranging from 0.25 to 5 mM (0.25, 0.5, 1, 2, 3, 4, and 5 mM). Michaelis-Menten analysis revealed distinct kinetic parameters, including V_max_, K_M_, k_cat_, and k_cat_/K_M_, under each sterol condition (Table 1, Fig. 3C). In the presence of ergosterol, the apparent V_max_, K_M_, and k_cat_ were 1505 ± 92 nmol min^-1^ mg^-1^, 1.82 ± 0.28 mM, and 3.96 ± 0.24 s^-1^, respectively. The V_max_ observed with ergosterol was more than twofold higher than that observed with CHS or cholesterol and approximately fivefold higher than in the absence of sterol. Similar increases were observed for k_cat_ under ergosterol-stimulated conditions. In contrast, ergosterol resulted in the highest apparent K_M_ value, corresponding to a 1.5–2.5-fold increase compared with CHS-, cholesterol-, or no-sterol conditions, indicating reduced apparent ATP affinity. Together, these results demonstrate that sterols differentially modulate ABCG5/G8 catalytic properties. Ergosterol primarily enhances ATP hydrolysis by increasing the maximal catalytic rate of the transporter.

**Table 1.**

| | $V_{\max}$<br>(nmol·min<sup>-1</sup>·mg<sup>-1</sup>) | $K_M$<br>(mM) | $k_{\text{cat}}$<br>(s<sup>-1</sup>) | $k_{\text{cat}}/K_M$<br>(10<sup>3</sup> M<sup>-1</sup>·s<sup>-1</sup>) | n* |
| --- | --- | --- | --- | --- | --- |
| **ATP-dependant ATPase activity** |  |  |  |  |  |
| **ABCG5<sub>WT</sub>/G8<sub>WT</sub> (Ergosterol)** | 1505 ± 92 | 1.82 ± 0.28 | 3.96 ± 0.24 | 2.18 ± 0.36 | 3 |
| **ABCG5<sub>WT</sub>/G8<sub>WT</sub> (CHS)** | 627 ± 39 | 1.10 ± 0.21 | 1.65 ± 0.10 | 1.50 ± 0.33 | 3 |
| **ABCG5<sub>WT</sub>/G8<sub>WT</sub> (Cholesterol)** | 693 ± 39 | 0.70 ± 0.15 | 1.83 ± 0.10 | 2.60 ± 0.56 | 3 |
| **ABCG5<sub>WT</sub>/G8<sub>WT</sub> (No sterol)** | 305 ± 34 | 0.88 ± 0.33 | 0.80 ± 0.09 | 0.91 ± 0.35 | 3 |
| **Sterol-dependant ATPase activity** |  |  |  |  |  |
| **ABCG5<sub>WT</sub>/G8<sub>WT</sub> (Ergosterol)** | 2819 ± 271 | 0.42 ± 0.07 | 7.43 ± 0.71 | 18 ± 3.40 | 3 |
| **ABCG5<sub>WT</sub>/G8<sub>WT</sub> (CHS)** | 888 ± 40 | 0.39 ± 0.04 | 2.34 ± 0.11 | 6.05 ± 0.73 | 3 |
| **ABCG5<sub>WT</sub>/G8<sub>WT</sub> (Cholesterol)** | 1395 ± 116 | 0.52 ± 0.09 | 3.67 ± 0.31 | 7.08 ± 1.41 | 3 |

\*n, number of independent assays.

Sterol concentration-dependent assays were also performed in the presence of 5 mM ATP and gradually increasing concentrations of ergosterol, cholesterol, or CHS (Table 1, Fig. 3D). For cholesterol and CHS, sterol concentrations of 0.25, 0.35, 0.5, 0.75, and 1 mM were tested. Due to the limited solubility of ergosterol under the assay conditions, particularly in the presence of soy PC, concentrations of 0.05, 0.1, 0.2, 0.25, 0.3, 0.35, and 0.4 mM were used instead. Under ergosterol-supplemented conditions, we observed a V_max_ of 2819 ± 271 nmol min^-1^ mg^-1^, a K_M_ of 0.42 ± 0.07 mM, and a k_cat_ of 7.43 ± 0.71 s^-1^. Compared to the kinetic parameters achieved in the presence of cholesterol or CHS, these values represent a 2–3-fold higher V_max_ and k_cat_, whereas KM values were comparable across conditions (fold difference of 1). The kinetic parameters obtained for CHS were consistent with our previous report (19).

## Discussion

In the present study, we combined cryo-EM structural determination, MD simulations, site-directed mutagenesis, and ATPase assays to gain mechanistic insight into sterol interactions with ABCG5/G8. Our cryo-EM structure revealed a sterol-like density located at the TMD interface between ABCG5 and ABCG8. Structural analysis indicated that this cavity is deeply buried within the TMDs and possesses a defined opening connecting the pocket to the surrounding membrane environment. This suggests that sterol molecules entering the pocket may be transiently stabilized prior to ATP-dependent conformational rearrangements. A comparable ligand-binding cavity has also been observed in ABCG2 structures (23).

To identify the ligand corresponding to the observed density, we compared the structural compatibility of different sterol species with the cryo-EM density. Structural fitting and refinement showed higher CC values for ergosterol, or cholesterol compared with CHS, indicating that the structural geometry and physicochemical properties of CHS are less compatible with the observed density. In addition, MD simulations showed that CHS was less stable in the pose corresponding to the observed density compared to cholesterol or ergosterol. Together, these results suggest that the observed cryo-EM density is most consistent with ergosterol.

Interestingly, a previous structural study reported a cholesterol molecule occupying this site after prolonged incubation with cholesterol (PDB ID: 7R8B) (24). These observations suggest that the cavity may accommodate multiple sterol species and that sterol occupancy may be influenced by the sterol composition and saturation state of the surrounding membrane environment. Therefore, aligned with our MD and structural analysis, the functional effects of different sterols on ABCG5/G8 activity may arise from their distinct interactions within the binding cavity. Comparison of the ergosterol-bound structure with the previously reported cholesterol-bound structure (PDB ID: 7R8B)(24), provided further insight into the molecular basis of differential sterol recognition within this cavity. The comparison revealed differences in residues involved in sterol binding and their subtle side chain rearrangements (Fig. S6). Specifically, ABCG8 asparagine 564 (ABCG8_N564_) was shown to contribute to cholesterol binding but does not participate in the corresponding ergosterol-binding interaction (Fig. S6B). Whether this residue contributes to sterol selectivity through differential interactions with cholesterol and ergosterol remains to be determined.

Despite the identification of ergosterol as the most likely ligand corresponding to the observed sterol density, direct evidence for an interaction between ergosterol and ABCG5/G8 had not previously been reported. Interestingly, previous studies in rats have shown that orally administered ergosterol is poorly retained, with the majority of the sterol excreted in feces and only minor amounts detected in circulation (25,26). These findings raise the possibility that intestinal sterol transport mechanisms may contribute to regulating ergosterol absorption and bioavailability. To assess whether ergosterol interacts with and modulates ABCG5/G8 activity, we measured ABCG5_WT_/G8_WT_ ATPase activity in the presence of ergosterol, cholesterol, and CHS applying an in vitro assay. ATPase assays revealed that all three sterols stimulated ABCG5/G8 ATPase activity, with ergosterol producing a markedly stronger stimulation compared with either cholesterol or CHS. Interestingly, cholesterol and CHS induced comparable levels of ATPase activation despite the distinct and bulkier headgroup of CHS, suggesting that the sterol hydroxyl group alone is not the determinant of transporter activation.

The differential activation of ABCG5/G8 by ergosterol compared with cholesterol and CHS encouraged us to investigate the structural determinants underlying this sterol-specific regulation. The unique positioning of ABCG5_Y432_ within the sterol-binding cavity suggested that this residue may represent a key determinant linking sterol recognition to transporter activation. ABCG5_Y432_ is located directly beneath the sterane ring of ergosterol within the binding cavity, and MD simulations identified this residue as one of the most favorable energetic contributors to sterol interactions (Fig. 2C). In addition, this residue forms part of two conserved structural features within ABCG5, the aromatic clamp and polar relay, which are believed to contribute to the structural organization and functional regulation of ABCG5/G8 (17,18). This position is conserved as phenylalanine across most ABCG transporters, including the cholesterol transporters ABCG1 and ABCG4, whereas ABCG5 uniquely contains a tyrosine residue at this site (17,27). The presence of a hydroxyl group at this conserved aromatic position may provide an additional interaction capability that contributes to the specialized sterol-sensing properties and functional regulation of ABCG5/G8. Functional analysis of the ABCG5_Y432F_/G8_WT_ protein further supported this hypothesis. While this substitution had minimal effects on cholesterol-stimulated ATPase activity, it substantially reduced ergosterol-dependent activation, suggesting that ABCG5_Y432_ contributes to the differential modulation of ABCG5/G8 activity by distinct sterol species within this cavity. Notably, a previous study demonstrated that alanine substitution of this residue severely impaired biliary cholesterol transport (18). Furthermore, we have previously postulated the differential sterol-transport mechanisms between ABCG1 and ABCG5/G8. This present study on the degenerative residue on ABCG5 further strengthen our hypothesis about substrate selectivity of ABCG sterol transporters by utilizing the conserved aromatic clamp structural motif (17).

Beyond the internal sterol-binding cavity identified in our cryo-EM structure, our previous crystal structure revealed an additional, distinct sterol interaction site at the surface-exposed interface between the ABCG5/G8 TMDs (17). In the crystal structure, the sterol density was observed at the ABCG8-ABCG5 transmembrane interface but was not detected at the opposite ABCG5-ABCG8 interface. Consistently, molecular docking analyses did not identify a favorable sterol-binding pose at the corresponding site on the ABCG5-facing side (17) (Fig. 4C). Structural analysis shows that residues ABCG5_I539_ and Leucine 543 (ABCG5_L543_) and ABCG8 Leucine 458 (ABCG8_L458_) and phenylalanine 461 (ABCG8_F461_) partition the transmembrane cavity into asymmetric subcavities. The cavity accessible from the ABCG5-ABCG8 interface is larger and can accommodate sterols, whereas the corresponding cavity on the ABCG8-ABCG5 side is substantially more restricted and may be less favorable for sterol entry (Fig. 4C). Together, these observations support a model in which sterols may access the internal cavity from the ABCG5-ABCG8 side, where they could be transiently accommodated before ATP-driven conformational changes promote transport (Fig. 4A). In contrast, sterols interacting with the ABCG8-ABCG5 side may preferentially associate with the surface-exposed site observed in the crystal structure, raising the possibility that this site contributes to sterol sensing or regulation rather than serving as a direct transport intermediate (Fig. 4C) (28). Together, these two structurally distinct sterol interaction sites suggest an asymmetric sterol-binding landscape within ABCG5/G8 (17).

**Figure 4.**
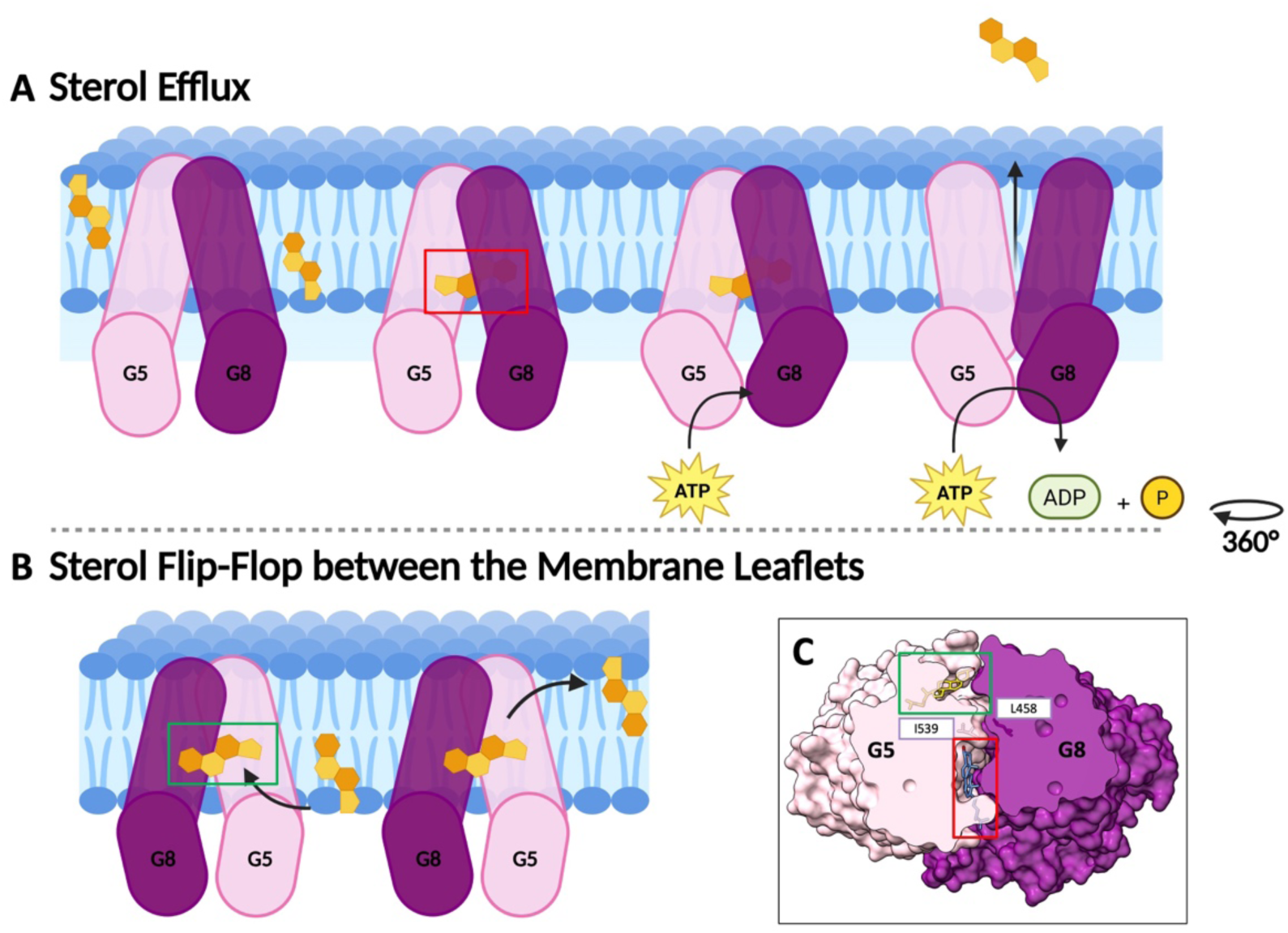
Proposed functional model of ABCG5/G8 highlighting sterol-binding sites identified in the crystal structure (PDB ID: 8CUB) (17) and the cryo-EM structure determined in this study. **A.** The model depicts sterol efflux through a binding site observed in the cryo-EM structure, with the sterol located between the TMDs indicated in the red box corresponding to ergosterol. Sterols are proposed to enter from the membrane into the binding cavity, where ATP binding and hydrolysis drive conformational changes that enable substrate release. **B.** Model of cholesterol translocation along the protein surface, illustrating a proposed flip-flop mechanism between membrane leaflets. The cholesterol molecule shown in the green box corresponds to the bound cholesterol observed in the crystal structure (PDB ID: 8CUB) (17). **C.** Surface representation of ABCG5/G8 (top view) showing sterol-binding sites identified in both structures. The site highlighted in red corresponds to that shown in panel A, while the site in green corresponds to panel B. Key residues, including ABCG5_I539_ and ABCG8_L458_, partition the transmembrane cavity into asymmetric subcavities. The ABCG5-ABCG8-facing cavity is spacious and accessible to sterols, whereas the ABCG8-ABCG5-facing cavity is more restricted and is not predicted to accommodate sterol entry.

An alternative hypothesis is that the surface-exposed binding site may represent an intermediate site during sterol translocation between the two membrane leaflets (Fig. 4B). This interpretation is consistent with molecular dynamics studies of P-glycoprotein, which have reported cholesterol “flip-flop” events along the protein surface, suggesting that surface-associated interactions may facilitate lateral sterol diffusion within the membrane (29). In cholesterol-enriched membrane environments, where local sterol abundance is elevated and lipid packing is altered, such surface-associated interactions may become more prevalent, potentially making this site responsive to changes in membrane sterol composition (29,30). Overall, these observations support a model in which surface-associated sterol binding may contribute to regulatory interactions, whereas the internal cavity likely represents the primary pathway for active sterol transport. Further biochemical and biophysical approaches are required to define the functional relationship between these sites and determine their broader implications for the transport mechanisms of homologous ABC transporters.

In conclusion, this study provides structural, computational, and functional evidence identifying a sterol-binding cavity within ABCG5/G8 and demonstrating that this transporter differentially responds to distinct sterol species. Furthermore, our findings highlight potential molecular determinants underlying sterol-specific regulation within ABCG sterol transporters. Our results further suggest that ABCG5/G8 may contribute to regulating the bioavailability of dietary ergosterol. Given the reported antioxidant and anti-inflammatory properties of ergosterol, its role as a precursor for vitamin D₂ biosynthesis in fungi, and growing interest in its therapeutic potential (3), ABCG5/G8-mediated regulation of ergosterol absorption and bioavailability warrants consideration during the development and optimization of orally administered ergosterol-based interventions.

## Material and Method

### Mutant construct generation and cloning

The ABCG5_Y432F_/G8_WT_ mutant construct used in this study was generated following our previously published protocol (19). Expression vectors (pLIC) carrying human ABCG5 (NCBI accession number NM_022436) and ABCG8 (NCBI accession number NM_022437) were previously generated in our laboratory from the pPICZB vector (Invitrogen). The pLIC-ABCG5 construct contained a 6His_Gly_6His affinity tag on ABCG5. The pLIC-ABCG8 construct contained a C-terminal rhinovirus 3C protease cleavage site followed by a calmodulin-binding peptide (CBP) tag. To generate the missense mutants in this study, we performed site-directed mutagenesis by using ABCG5_WT_ as the template. The polymerase chain reaction (PCR) primers (for ABCG5_Y432F_) used in this study (Forward primer sequence: 5’ caccccgttcacaggcatgctg 3’, Reverse primer sequence: 5’ cagcatgcctgtgaacggggtg 3’) were designed by Benchling (31) and ordered from Thermo Fisher Scientific. The ABCG5_WT_/ABCG8_G216D_ mutant was generated previously (19).

The PCR was performed using a reaction master mixture containing 1 ug of DNA template (ABCG5_WT_), 5 uM of each forward and reverse primers, 1 ul DMSO, 10 ul 10x GC buffer, 1 ul DNTs, 0.5 ul Phusion enzyme (all from NEB) and ddH2O. The thermal cycle program for PCR consisted of 35 cycles, and each cycle comprised 95°C 1 min, 56°C 30 s, 72°C for 8 min. PCR products were detected by a 0.7% agarose (Bio-Rad Laboratories) gel electrophoresis.

PCR products were treated for 1 hour with DpnI (NEB) at 37°C and analyzed by 0.7% agarose gel electrophoresis. Mutant plasmids were subsequently transformed into XL1-Blue competent *Escherichia coli* cells using a heat-shock protocol according to the manufacturer’s instructions (Novagen/Agilent Technologies). Colonies were selected on LB agar plates containing 25 ug/ml Zeocin (Invitrogen/Thermo Fisher Scientific). Some colonies were selected for plasmid minipreparation using the standard alkaline lysis method as described in Molecular Cloning protocol by Sambrook and Russell (32). Purified plasmid DNA was sequenced at StemCore Laboratories, the genomics core facility in Ottawa.

Sequence-verified mutant pLIC-ABCG5_Y432F_ and pLIC-ABCG8_WT_ plasmids were linearized by PmeI (NEB) digestion and ethanol purified. 5-10 ug of linearized pLIC-ABCG5_Y432F_ and pLIC-ABCG8_WT_ plasmids were transformed into freshly prepared KM71H *Pichia pastoris* competent cells by electroporation. Transformed yeast cells were plated on Yeast Extract Peptone Dextrose Sorbitol (YPDS) agar plates containing various concentrations of Zeocin and incubated for 3–10 days. The largest colonies were selected and screened for protein expression. Protein expression of both ABCG5 and ABCG8 was evaluated by western blot analysis using anti-His tag and anti-ABCG8 antibodies (Invitrogen) respectively. (Clarity Western enhanced chemiluminescence (ECL) substrate used for western blot detection, and 30% acrylamide and ammonium persulfate used for sodium dodecyl sulfate-polyacrylamide gel electrophoresis (SDS-PAGE) gel casting were obtained from Bio-Rad Laboratories).

### Cell Culture, Microsomal Membrane Preparation, and Protein Purification

Cell growth and membrane preparation were performed according to our protocol published previously (17). In brief, yeast cells were grown in 6-L cultures in minimal glycerol yeast nitrogen base (MGY) and were harvested in sucrose lysis buffer and broken using a bead beater to prepare approximately 100 mL of microsomal membranes.

Protein purification was performed according to the previously published protocol (17) with minor modifications, as described below. Membranes were solubilized in an equal volume of solubilization buffer containing 50 mM Tris-HCl (pH 8.0), 100 mM NaCl (Bioshop), 10% glycerol (Bioshop), 2% (w/v) β-dodecyl maltoside (β-DDM; Anatrace), 0.5% (w/v) cholate (MilliporeSigma), 0.25% (w/v) cholesteryl hemisuccinate Tris (CHS-Tris; Anatrace), 5 mM imidazole (Bioshop), 5 mM β-mercaptoethanol (β-ME) (MilliporeSigma), 2 μg/mL leupeptin, 2 μg/mL pepstatin A, and 2 mM PMSF (all from Bioshop) for 2 hours at 4 °C. Following ultracentrifugation at 30,000 rpm for 30 min, the clarified supernatant containing solubilized membrane proteins was incubated overnight with 20 mL nickel-nitrilotriacetic acid (Ni-NTA) affinity resin (Qiagen). The resin was subsequently packed into a column and washed with 10 column volumes of Buffer B containing 50 mM Tris-HCl (pH 7.5), 100 mM NaCl, 0.05% (w/v) cholate, 0.1% (w/v) β-DDM, 0.01% (w/v) CHS, and 25 mM imidazole. Bound proteins were eluted using Buffer B supplemented with 250 mM imidazole. Peak fractions were identified using a colorimetric protein assay using Bradford reagent (Bioshop) and subsequently pooled.

In this study, two ABCG8 constructs were used: an untagged construct, and a construct containing a C-terminal CBP tag. Samples containing CBP-tagged ABCG8 were subjected to a second affinity purification step using a 5-mL calmodulin affinity column (Agilent Technologies) when further purification was required. For this purpose, pooled Ni-NTA fractions were diluted 1:1 with CBP wash buffer containing 50 mM Tris-HCl (pH 7.5), 100 mM NaCl, 0.1% (w/v) β-DDM, 0.05% (w/v) cholate, 0.01% (w/v) CHS, 1 mM CaCl_2_, and 1 mM MgCl_2_ (Bioshop), and loaded onto the CBP column. Following washing, ABCG5/G8 was eluted using buffer containing 50 mM Tris-HCl (pH 7.5), 300 mM NaCl, 2 mM EGTA (MilliporeSigma), 0.05% (w/v) cholate, and 0.01% (w/v) CHS.

All constructs were subsequently concentrated using 100-kDa molecular weight cutoff filters (Cytiva) and further purified by size-exclusion chromatography on a Superdex 200 10/300 GL column using an ÄKTA fast protein liquid chromatography (FPLC) system. The gel filtration buffer consisted of 10 mM Tris-HCl (pH 7.5), 100 mM NaCl, 0.1% (w/v) DDM, 0.05% (w/v) cholate, and 0.01% (w/v) CHS. Peak fractions were pooled and concentrated using 100-kDa molecular weight cutoff filters. Protein concentration was estimated by SDS-PAGE using bovine serum albumin (BSA) (MilliporeSigma) standards. Samples were stored for subsequent ATPase assays and cryo-EM analysis.

### Cryo-EM grid preparation, data collection, and processing

For grid preparation, 4 μL of purified ABCG5/ABCG8 (∼1.5 mg/mL) which was incubated overnight with 2 mM CHS and supplemented with 10 mM AMP-PNP (MilliporeSigma) and MgCl₂ was applied to glow-discharged Quantifoil R1.2/1.3 300-mesh copper grids (Quantifoil Micro Tools GmbH). Grids were blotted for 3s at 4 °C and 100% relative humidity and flash-frozen in liquid ethane using a FEI Vitrobot. Data were collected on a Titan Krios transmission electron microscope (Thermo Fisher Scientific) operated at 300 kV and equipped with a BioQuantum energy filter (Gatan) and a K3 direct electron detector (Gatan) at the Cryo-EM facility of Academia Sinica (Taipei, Taiwan). Dose-fractionated movies were recorded in super-resolution mode at a nominal magnification of ×105,000, corresponding to a physical pixel size of 0.84 Å. Six-second exposures were fractionated into 0.2-s frames. The total accumulated dose was ∼50 electrons/Å².

Two datasets comprising a total of 21,287 micrographs were collected for further processing. Motion correction was performed applying MotionCor2 on-site (33). All subsequent image processing was performed in CryoSPARC (34). Contrast transfer function (CTF) parameters were estimated using patch CTF estimation (35). Particles were initially picked using TOPAZ and blob picking to generate reference-free 2D class averages (34,36). The best 2D classes were selected as templates for iterative particle picking. Approximately 3 million particles were extracted and subjected to several rounds of 2D classification to remove junk particles. A final subset of 455,323 particles was selected and subjected to heterogeneous refinement, followed by additional heterogeneous, homogeneous, non-uniform, and local refinement to improve map quality (34,37). The final 3D reconstruction reached an overall resolution of 3.56 Å, as determined by the gold-standard FSC 0.143 criterion (38).

Model building was performed by real-space refinement in Phenix using our previously published crystal structure (PDB ID: 8CUB) as an initial model. Manual model adjustment and inspection of electron density were performed using COOT (39). Despite the application of AMP-PNP during cryo-EM sample preparation, no additional density was observed in the nucleotide-binding sites (NBSs). Structural figures were prepared using PyMOL and UCSF Chimera (40,41).

### Software used in this project for cryo-EM data processing was curated by SBGrid (42)

#### ATPase assay performance and data processing

ATPase activity of purified ABCG5/G8 and its mutants was measured using a colorimetric inorganic phosphate detection assay that we reported in our previous publication (19). The protocol with some modifications is as follow: 1 µg of protein was used per reaction with 2 mM DTT. Lipids and sterols dissolved in chloroform were dried under nitrogen gas in glass tubes and vacuum-desiccated overnight. Dried lipids (2 mg/mL) were resuspended in assay buffer containing 10% glycerol, 50 mM Tris pH 7.5, 100 mM NaCl, 1.5% sodium cholate, and 0.1% DDM, then heated and sonicated until clear.

Reactions were assembled at room temperature by mixing protein, DTT, lipid mixture (2 mg/mL soy PC) (Avanti) and varying concentrations of sterols), and double-distilled water. Reaction mixtures were subsequently incubated at 37°C and initiated by the addition of 5 mM ATP (or different concentrations of ATP), 7.5 mM MgCl_2_, and 10 mM NaN_3_. Reactions were quenched at designated time points (0–20 min) by transferring 8 μL of reaction mixture into 25 μL of ice-cold 5% SDS in 5 mM HCl in a 96-well plate. Inorganic phosphate standards (0–200 μM) were prepared in parallel using 50 mM Tris-HCl buffer (pH 7.5). Released inorganic phosphate was detected by the addition of 50 μL freshly prepared solution containing 1% ammonium molybdate (Bioshop) and 1.5% ascorbic acid in HCl (MilliporeSigma), followed by incubation on ice for 10 min. Subsequently, 75 μL of a freshly prepared solution containing bismuth citrate and sodium citrate (both from Millipore Sigma) in HCl was added, and samples were incubated at 37°C for an additional 10 min. Absorbance at 695 nm was measured using a plate reader. GraphPad Prism 11 was used to perform nonlinear regression and ordinary one-way ANOVA, with a p-value of ≤0.05 considered significant from three independent experiments. The kinetic parameters were calculated by nonlinear Michaelis-Menten curve fitting using GraphPad Prism 11 (43).

#### MD simulations

The ABCG5/G8 structure (PDB ID: 8CUB) was retrieved from the protein data bank with missing residues modeled using MODELLER (44). All sterols were docked using AutoDock VINA (45). The docking poses that best matched the experimentally resolved sterol orientations were selected for further analysis. The CHARMM-GUI web server was used to prepare molecular dynamics systems, using the recommended protonation states (46,47). The Orientations of Proteins in Membranes (OPM) database was used to orient the protein within a 118 Å × 118 Å membrane patch composed of POPC and cholesterol at a 5:1 molar ratio (48). A 13 Å water layer of TIP3P water molecules was added above and below the membrane, followed by ionization with 0.15 M NaCl. Hydrogen mass repartitioning was applied to allow for more efficient computational sampling using 4 fs time steps (49).

All-atom molecular dynamics simulations were performed using NAMD3 (50) with the CHARMM36 and CHARMM36m forcefields (51,52). Systems underwent a three-step minimization and equilibration protocol. Initially, the membrane was relaxed for 10 ns, followed by an additional 10 ns relaxation of the membrane and solvent components. Lastly, the backbone atoms were restrained while the remaining components were allowed to relax for 40 ns. This was followed by 225 ns of independent production simulations.

Short-range nonbonded interactions were truncated at 12 Å, with a smoothing function applied beginning at 10 Å. Long-range electrostatic interactions were treated using the Particle Mesh Ewald (PME) method (53) with a grid density of 1 Å⁻¹. Water molecules were constrained using the SETTLE algorithm (54), while bonds involving hydrogen atoms were constrained using the RATTLE/SHAKE algorithms (55). Pressure and temperature were maintained at 1 atm and 310 K, respectively, using a Langevin piston barostat and a Langevin thermostat.

RMSD analysis was done using MDAnalysis (56). Per-residue decomposition of binding energy was done using AmberTools23 (57). NAMD topology files were converted to AMBER topology using the ParmEd tool. Molecular Mechanics Generalized Born Surface Area (MM-GBSA) was used to approximate the binding energy (58). Relevant residues were selected if they averaged ≤ −0.5 or ≥ +0.5 kcal/mol contribution to ligand binding in any trajectory. The first 25 ns of each simulation were discarded to allow for stabilization of the protein backbone. All analyses were performed using the remaining 200 ns of each trajectory.

## Data Availability

The cryo-EM data have been deposited in PDB and EMDB under the accession codes 37WK and EMD-78562, respectively.

## Acknowledgement

This work was supported by a startup grant from the University of Ottawa, a National New Investigator Award from the Heart and Stroke Foundation of Canada, and a Canadian Institutes of Health Research Project Grant (PJT-180640) to JYL, Academia Sinica intramural fund, an Academia Sinica Investigator Award (AS-IV-114-L04) to STDH, an Academia Sinica – University of Ottawa Travel Support for Research Projects to STDH and JYL, and a Mitacs Globalink Research Award (GRA, IT47640) and an Academia Sinica Taiwan International Graduate Program X (TIGP-X) to DF, JYL and STDH. DF is a recipient of a CIHR Canada Graduate Research Scholarship-Doctoral (CGRS-D). Part of this work were used to fulfill in part the requirement for the degrees of Honours Bachelor of Science (QFX).

We thank the Academia Sinica Cryo-EM Center (AS-CFII-111-210) for data collection, and the Distributed Cloud Operating System (DiCOS) of the Academia Sinica Grid Computing (ASGC), both are funded by the Academia Sinica Core Facility and Innovative Instrument Project. We also thank the StemCore Laboratories, the Genomics Core Facility at the Ottawa Hospital Research Institute, for DNA sequencing services.

We are grateful to Ms. Gonca Gursu and Dr. Jean-François Couture for their valuable feedback on the manuscript and to Mr. Barry Cao and Ms. Kimia Ebrahimi for their technical assistance.

ChatGPT (OpenAI, GPT-5.5) was utilised to assist with improving the grammar, clarity, and readability of the manuscript.

## Statement of Author Contributions

Conceptualization: FR, JYL

Data curation: FR, IFO, DF, ZWW, KVS, QFX, YCC

Formal analysis: FR, IFO, DF, ZWW

Funding acquisition: STDH, JYL

Investigation: FR, IFO, DF, ZWW

Project administration: JYL

Resources: YCC, STDH, JYL

Supervision: STDH, JYL

Validation: FR, IFO, DF, STDH, JYL

Visualization: FR, IFO, DF

Writing-original draft: FR, IFO, DF, JYL

Writing – review & editing: FR, IFO, DF, ZWW, KVS, QFX, YCC, STDH, JYL

## References

1. Hartmann MA. Plant sterols and the membrane environment. Trends Plant Sci. 1998 May 1;3(5):170–5. doi:10.1016/S1360-1385(98)01233-3

2. Zio S, Tarnagda B, Tapsoba F, Zongo C, Savadogo A. Health interest of cholesterol and phytosterols and their contribution to one health approach: Review. Heliyon. 2024 Nov 15;10(21):e40132. doi:10.1016/J.HELIYON.2024.E40132 PubMed PMID: 39583830.

3. Rangsinth P, Sharika R, Pattarachotanant N, Duangjan C, Wongwan C, Sillapachaiyaporn C, et al. Potential Beneficial Effects and Pharmacological Properties of Ergosterol, a Common Bioactive Compound in Edible Mushrooms. Foods 2023, Vol 12, Page 2529. 2023 Jun 29;12(13):2529. doi:10.3390/FOODS12132529

4. Zio S, Tarnagda B, Tapsoba F, Zongo C, Savadogo A. Health interest of cholesterol and phytosterols and their contribution to one health approach: Review. Heliyon. 2024 Nov 15;10(21):e40132. doi:10.1016/J.HELIYON.2024.E40132 PubMed PMID: 39583830.

5. Mau JL, Chen PR, Yang JH. Ultraviolet Irradiation Increased Vitamin D2 Content in Edible Mushrooms. J Agric Food Chem. 1998;46(12):5269–72. doi:10.1021/JF980602Q

6. McNamara DJ. Dietary cholesterol and atherosclerosis. Biochimica et Biophysica Acta (BBA) - Molecular and Cell Biology of Lipids. 2000 Dec 15;1529(1–3):310–20. doi:10.1016/S1388-1981(00)00156-6 PubMed PMID: 11111098.

7. Schade DS, Shey L, Eaton RP. Cholesterol Review: A Metabolically Important Molecule. Endocrine Practice. 2020 Dec 1;26(12):1514–23. doi:10.4158/EP-2020-0347 PubMed PMID: 33471744.

8. Bloch K. The biological synthesis of cholesterol. Science (1979). 1965 Oct 1;150(3692):19–28. doi:10.1126/SCIENCE.150.3692.19/ASSET/2D2B83E1-7C63-456D-835E-B11140E12B71/ASSETS/SCIENCE.150.3692.19.FP.PNG PubMed PMID: 5319508.

9. Garçon D, Berger JM, Cariou B, Le May C. Transintestinal cholesterol excretion in health and disease. Curr Atheroscler Rep. 2022 Mar 1;24(3):153–60. doi:10.1007/S11883-022-00995-Y PubMed PMID: 35138569.

10. Heinemann T, Axtmann G, Bergmann K Von. Comparison of intestinal absorption of cholesterol with different plant sterols in man*. Eur J Clin Invest. 1993 Dec 1;23(12):827–31. doi:10.1111/J.1365-2362.1993.TB00737.X PubMed PMID: 8143759.

11. Tabas I. Consequences of cellular cholesterol accumulation: basic concepts and physiological implications. J Clin Invest. 2002 Oct 1;110(7):905–11. doi:10.1172/JCI16452 PubMed PMID: 12370266.

12. Klett EL, Lee MH, Adams DB, Chavin KD, Patel SB. Localization of ABCG5 and ABCG8 proteins in human liver, gall bladder and intestine. BMC Gastroenterol. 2004 Sep 21;4(1):21-. doi:10.1186/1471-230X-4-21/FIGURES/6 PubMed PMID: 15383151.

13. Wang J, Mitsche MA, Lütjohann D, Cohen JC, Xie XS, Hobbs HH. Relative roles of ABCG5/ABCG8 in liver and intestine. J Lipid Res. 2015 Feb 1;56(2):319–30. doi:10.1194/jlr.M054544 PubMed PMID: 25378657.

14. Bhattacharyya AK, Connor WE. β-Sitosterolemia and Xanthomatosis: A NEWLY DESCRIBED LIPID STORAGE DISEASE IN TWO SISTERS. J Clin Invest. 1974 Apr 1;53(4):1033–43. doi:10.1172/JCI107640 PubMed PMID: 4360855.

15. Salen G, Horak I, Rothkopf M, Cohen JL, Speck J, Tint GS, et al. Lethal atherosclerosis associated with abnormal plasma and tissue sterol composition in sitosterolemia with xanthomatosis. J Lipid Res. 1985 Sep 1;26(9):1126–33. doi:10.1016/S0022-2275(20)34286-3 PubMed PMID: 4067433.

16. Miettinen TA. Phytosterolaemia, xanthomatosis and premature atherosclerotic arterial disease: a case with high plant sterol absorption, impaired sterol elimination and low cholesterol synthesis. Eur J Clin Invest. 1980 Feb 1;10(1):27–35. doi:10.1111/J.1365-2362.1980.TB00006.X PubMed PMID: 6768564.

17. Farhat D, Rezaei F, Ristovski M, Yang Y, Stancescu A, Dzimkova L, et al. Structural Analysis of Cholesterol Binding and Sterol Selectivity by ABCG5/G8. J Mol Biol. 2022 Oct 30;434(20):167795. doi:10.1016/J.JMB.2022.167795 PubMed PMID: 35988751.

18. Lee JY, Kinch LN, Borek DM, Wang J, Wang J, Urbatsch IL, et al. Crystal structure of the human sterol transporter ABCG5/ABCG8. Nature 2016 533:7604. 2016 May 4;533(7604):561–4. doi:10.1038/nature17666 PubMed PMID: 27144356.

19. Xavier BM, Zein AA, Venes A, Wang J, Lee JY. Transmembrane Polar Relay Drives the Allosteric Regulation for ABCG5/G8 Sterol Transporter. International Journal of Molecular Sciences 2020, Vol 21, Page 8747. 2020 Nov 19;21(22):8747. doi:10.3390/IJMS21228747 PubMed PMID: 33228147.

20. Wang Z, Stalcup LD, Harvey BJ, Weber J, Chloupkova M, Dumont ME, et al. Purification and ATP hydrolysis of the putative cholesterol transporters ABCG5 and ABCG8. Biochemistry. 2006 Aug 15;45(32):9929–39. doi:10.1021/BI0608055/SUPPL_FILE/BI0608055SI20060422_023206.PDF PubMed PMID: 16893193.

21. Sun Y, Wang J, Long T, Qi X, Donnelly L, Elghobashi-Meinhardt N, et al. Molecular basis of cholesterol efflux via ABCG subfamily transporters. Proc Natl Acad Sci U S A. 2021 Aug 24;118(34):e2110483118. doi:10.1073/PNAS.2110483118/SUPPL_FILE/PNAS.2110483118.SM01.MPG PubMed PMID: 34404721.

22. Zhang H, Huang CS, Yu X, Lee J, Vaish A, Chen Q, et al. Cryo-EM structure of ABCG5/G8 in complex with modulating antibodies. Communications Biology 2021 4:1. 2021 May 5;4(1):526-. doi:10.1038/s42003-021-02039-8 PubMed PMID: 33953337.

23. Orlando BJ, Liao M. ABCG2 transports anticancer drugs via a closed-to-open switch. Nature Communications 2020 11:1. 2020 May 8;11(1):2264-. doi:10.1038/s41467-020-16155-2 PubMed PMID: 32385283.

24. Sun Y, Wang J, Long T, Qi X, Donnelly L, Elghobashi-Meinhardt N, et al. Molecular basis of cholesterol efflux via ABCG subfamily transporters. Proc Natl Acad Sci U S A. 2021 Aug 24;118(34):e2110483118. doi:10.1073/PNAS.2110483118/SUPPL_FILE/PNAS.2110483118.SM01.MPG PubMed PMID: 34404721.

25. Tsugawa N, Okano T, Takeuchi A, Kayam M, Kobayashi T. Metabolism of Orally Administered Ergosterol and 7-Dehydrocholesterol in Rats and Lack of Evidence for Their Vitamin D Biological Activity. J Nutr Sci Vitaminol (Tokyo). 1992;38(1):15–25. doi:10.3177/JNSV.38.15 PubMed PMID: 1629783.

26. Zhao YY, Cheng XL, Liu R, Ho CC, Wei F, Yan SH, et al. Pharmacokinetics of ergosterol in rats using rapid resolution liquid chromatography–atmospheric pressure chemical ionization multi-stage tandem mass spectrometry and rapid resolution liquid chromatography/tandem mass spectrometry. Journal of Chromatography B. 2011 Jul 1;879(21):1945–53. doi:10.1016/J.JCHROMB.2011.05.025 PubMed PMID: 21664883.

27. Rezaei F, Farhat D, Gursu G, Samnani S, Lee JY. Snapshots of ABCG1 and ABCG5/G8: A Sterol’s Journey to Cross the Cellular Membranes. Int J Mol Sci. 2023 Jan 1;24(1):484. doi:10.3390/IJMS24010484/S1

28. Sharpe LJ, Rao G, Jones PM, Glancey E, Aleidi SM, George AM, et al. Cholesterol sensing by the ABCG1 lipid transporter: Requirement of a CRAC motif in the final transmembrane domain. Biochimica et Biophysica Acta (BBA) - Molecular and Cell Biology of Lipids. 2015 Jul 1;1851(7):956–64. doi:10.1016/J.BBALIP.2015.02.016 PubMed PMID: 25732853.

29. Thangapandian S, Kapoor K, Tajkhorshid E. Probing cholesterol binding and translocation in P-glycoprotein. Biochimica et Biophysica Acta (BBA) - Biomembranes. 2020 Jan 1;1862(1):183090. doi:10.1016/J.BBAMEM.2019.183090 PubMed PMID: 31676371.

30. Aussenac F, Tavares M, Dufourc EJ. Cholesterol Dynamics in Membranes of Raft Composition: A Molecular Point of View from 2H and 31P Solid-State NMR. Biochemistry. 2003 Feb 18;42(6):1383–90. doi:10.1021/BI026717B

31. Benchling [Biology Software]. (2024). Retrieved from https://benchling.com.

32. Molecular Cloning: A Laboratory Manual - Joseph Sambrook, David William Russell - Google Books [Internet]. [cited 2026 Jul 3]. Available from: https://books.google.ca/books?id=YTxKwWUiBeUC&printsec=frontcover&redir_esc=y#v=onepage&q&f=false

33. Li X, Mooney P, Zheng S, Booth CR, Braunfeld MB, Gubbens S, et al. Electron counting and beam-induced motion correction enable near-atomic-resolution single-particle cryo-EM. Nature Methods 2013 10:6. 2013 May 5;10(6):584–90. doi:10.1038/nmeth.2472 PubMed PMID: 23644547.

34. Punjani A, Rubinstein JL, Fleet DJ, Brubaker MA. cryoSPARC: algorithms for rapid unsupervised cryo-EM structure determination. Nature Methods 2017 14:3. 2017 Feb 6;14(3):290–6. doi:10.1038/nmeth.4169 PubMed PMID: 28165473.

35. Zivanov J, Nakane T, Scheres SHW. Estimation of high-order aberrations and anisotropic magnification from cryo-EM data sets in RELION-3.1. urn:issn:2052-2525. 2020 Feb 11;7(2):253–67. doi:10.1107/S2052252520000081

36. Bepler T, Morin A, Rapp M, Brasch J, Shapiro L, Noble AJ, et al. Positive-unlabeled convolutional neural networks for particle picking in cryo-electron micrographs. Nature Methods 2019 16:11. 2019 Oct 7;16(11):1153–60. doi:10.1038/s41592-019-0575-8 PubMed PMID: 31591578.

37. Punjani A, Zhang H, Fleet DJ. Non-uniform refinement: adaptive regularization improves single-particle cryo-EM reconstruction. Nature Methods 2020 17:12. 2020 Nov 30;17(12):1214–21. doi:10.1038/s41592-020-00990-8 PubMed PMID: 33257830.

38. Harauz G van HM. Exact filters for general geometry three dimensional reconstruction. Optik.. 1986 Feb;73(4):146–56.

39. Emsley P, Cowtan K. Coot: model-building tools for molecular graphics. urn:issn:0907-4449. 2004 Nov 26;60(12):2126–32. doi:10.1107/S0907444904019158 PubMed PMID: 15572765.

40. Pettersen EF, Goddard TD, Huang CC, Couch GS, Greenblatt DM, Meng EC, et al. UCSF Chimera—A visualization system for exploratory research and analysis. J Comput Chem. 2004 Oct 1;25(13):1605–12. doi:10.1002/JCC.20084 PubMed PMID: 15264254.

41. The PyMOL Molecular Graphics System, Version 3.0 Schrödinger, LLC.

42. Morin A, Eisenbraun B, Key J, Sanschagrin PC, Timony MA, Ottaviano M, et al. Collaboration gets the most out of software. Elife. 2013 Sep 10;2013(2). doi:10.7554/ELIFE.01456 PubMed PMID: 24040512.

43. Confidence intervals of proportions were calculated using the GraphPad QuickCalcs Web site: http://www.graphpad.com/quickcalcs/ConfInterval1.cfm (accessed 2025-2026).

44. Eswar N, Eramian D, Webb B, Shen MY, Sali A. Protein structure modeling with MODELLER. Methods Mol Biol. 2008;426:145–59. doi:10.1007/978-1-60327-058-8_8/FIGURES/4_8 PubMed PMID: 18542861.

45. Trott O, Olson AJ. AutoDock Vina: Improving the speed and accuracy of docking with a new scoring function, efficient optimization, and multithreading. J Comput Chem. 2010 Jan 30;31(2):455–61. doi:10.1002/JCC.21334 PubMed PMID: 19499576.

46. Lee J, Cheng X, Jo S, MacKerell AD, Klauda JB, Im W. CHARMM-GUI Input Generator for NAMD, Gromacs, Amber, Openmm, and CHARMM/OpenMM Simulations using the CHARMM36 Additive Force Field. Biophys J. 2016 Feb 16;110(3):641a. doi:10.1016/j.bpj.2015.11.3431

47. Wu EL, Cheng X, Jo S, Rui H, Song KC, Dávila-Contreras EM, et al. CHARMM-GUI Membrane Builder toward realistic biological membrane simulations. J Comput Chem. 2014 Oct 15;35(27):1997–2004. doi:10.1002/JCC.23702 PubMed PMID: 25130509.

48. Lomize MA, Pogozheva ID, Joo H, Mosberg HI, Lomize AL. OPM database and PPM web server: resources for positioning of proteins in membranes. Nucleic Acids Res. 2012 Jan 1;40(D1):D370–6. doi:10.1093/NAR/GKR703 PubMed PMID: 21890895.

49. Gao Y, Lee J, Smith IPS, Lee H, Kim S, Qi Y, et al. CHARMM-GUI Supports Hydrogen Mass Repartitioning and Different Protonation States of Phosphates in Lipopolysaccharides. J Chem Inf Model. 2021 Feb 22;61(2):831–9. doi:10.1021/ACS.JCIM.0C01360/ASSET/IMAGES/MEDIUM/CI0C01360_M002.GIF PubMed PMID: 33442985.

50. Phillips JC, Hardy DJ, Maia JDC, Stone JE, Ribeiro J V., Bernardi RC, et al. Scalable molecular dynamics on CPU and GPU architectures with NAMD. Journal of Chemical Physics. 2020 Jul 28;153(4). doi:10.1063/5.0014475/1064953 PubMed PMID: 32752662.

51. Huang J, Rauscher S, Nawrocki G, Ran T, Feig M, De Groot BL, et al. CHARMM36m: an improved force field for folded and intrinsically disordered proteins. Nature Methods 2016 14:1. 2016 Nov 7;14(1):71–3. doi:10.1038/nmeth.4067 PubMed PMID: 27819658.

52. Huang J, Mackerell AD. CHARMM36 all-atom additive protein force field: Validation based on comparison to NMR data. J Comput Chem. 2013 Sep 30;34(25):2135–45. doi:10.1002/JCC.23354 PubMed PMID: 23832629.

53. Essmann U, Perera L, Berkowitz ML, Darden T, Lee H, Pedersen LG. A smooth particle mesh Ewald method. J Chem Phys. 1995 Nov 15;103(19):8577–93. doi:10.1063/1.470117

54. Miyamoto S, Kollman PA. Settle: An analytical version of the SHAKE and RATTLE algorithm for rigid water models. J Comput Chem. 1992 Oct 1;13(8):952–62. doi:10.1002/JCC.540130805

55. Andersen HC. Rattle: A “velocity” version of the shake algorithm for molecular dynamics calculations. J Comput Phys. 1983 Oct 1;52(1):24–34. doi:10.1016/0021-9991(83)90014-1

56. Michaud-Agrawal N, Denning EJ, Woolf TB, Beckstein O. MDAnalysis: A toolkit for the analysis of molecular dynamics simulations. J Comput Chem. 2011 Jul 30;32(10):2319–27. doi:10.1002/JCC.21787 PubMed PMID: 21500218.

57. Case DA, Aktulga HM, Belfon K, Cerutti DS, Cisneros GA, Cruzeiro VWD, et al. AmberTools. J Chem Inf Model. 2023 Oct 23;63(20):6183. doi:10.1021/ACS.JCIM.3C01153 PubMed PMID: 37805934.

58. Genheden S, Ryde U. The MM/PBSA and MM/GBSA methods to estimate ligand-binding affinities. Expert Opin Drug Discov. 2015 May 1;10(5):449–61. doi:10.1517/17460441.2015.1032936 PubMed PMID: 25835573.

