## Supplementary Information for "A Sterol-Binding Cavity Underlies Sterol Recognition and Differential Activation of the ABCG5/G8"

**Supplementary Tables**

| Table S1 | |
| --- | --- |
| Data Collection and Processing Parameters | |
| Magnification | 105,000 |
| Voltage (kV) | 300 kV |
| Electron exposure (e–/Å2) | 1.05748 |
| Pixel size (Å) | 0.84 |
| Final Map Parameters | |
| Symmetry imposed | C1 |
| Initial images (no.) | 21,287 |
| Final particle (no.) | 227,299 |
| Model resolution (Å) | 3.56 |
| FSC threshold | 0.143 |
| Final Model Parameters | |
| Reference model | PDB ID: 8CUB |
| Chains | 2 |
| Protein residues | 1169 |
| Bonds (RMSD) | |
| Length (Å) (# > 4σ) | 0.004 (0) |
| Angles (°) (# > 4σ) | 0.604 (0) |
| MolProbity score | 1.75 |
| Clash score | 7.92 |
| Ramachandran plot (%) | |
| Outliers | 0.00 |
| Allowed | 3.72 |
| Favored | 96.28 |
| Rama-Z (Ramachandran plot Z-score, RMSD) | |
| whole (N = 1155) | 1.39 (0.26) |
| helix (N = 678) | 2.25 (0.20) |
| sheet (N = 84) | -0.69 (0.65) |
| loop (N = 393) | -1.15 (0.31) |
| Rotamer outliers (%) | 1.21 |
| CaBLAM outliers (%) | 2.89 |
| Ligand Parameters | |
| Ligand-#No | Ergosterol-1 |
| Mean CC for ligands | 0.80 |

**Supplementary Figures**

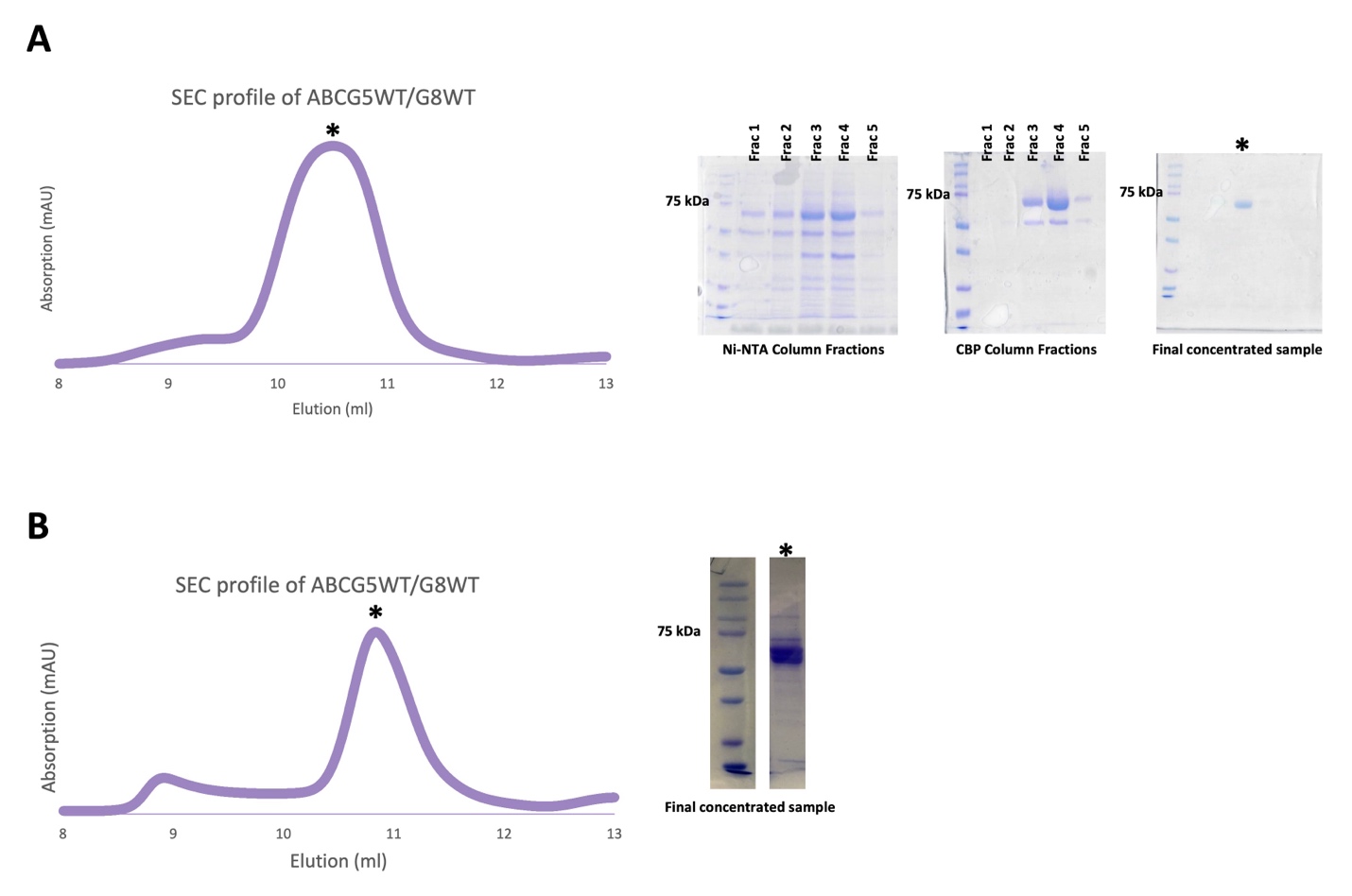

**Figure S1. Purification profiles of ABCG5_WT_/G8_WT_ constructs.** The ABCG5 construct was identical in all samples and contained a 6His-Gly-6His tag. ABCG8 contained a CBP tag in (A), no tag in (B).

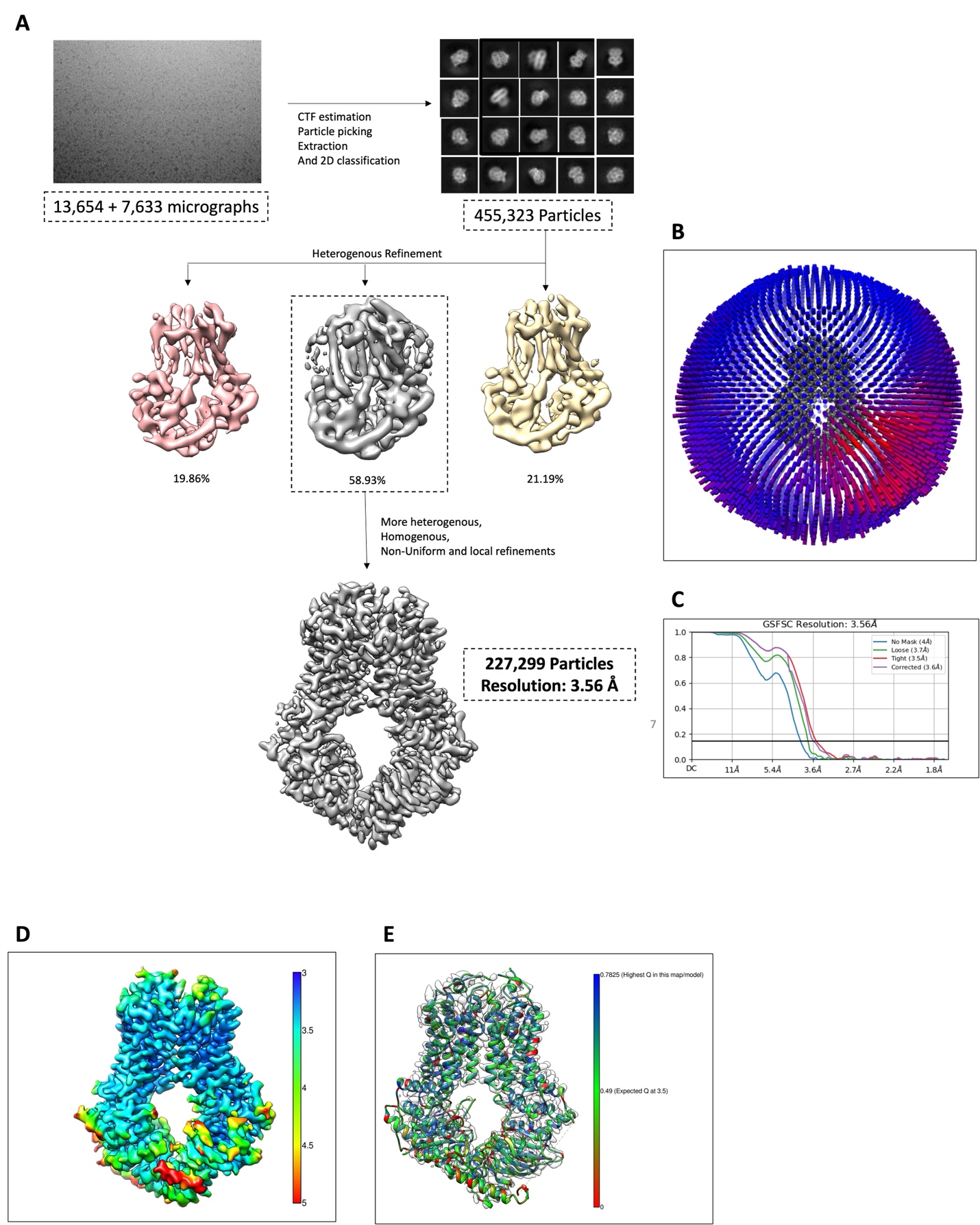

**Figure S2. Cryo-EM sample preparation and data processing. A.** Workflow of cryo-EM sample preparation and data processing. Representative motion-corrected micrographs (21,287 total) were subjected to CTF estimation, particle picking, and extraction, followed by multiple rounds of 2D classification. Selected particles (455,323) underwent heterogeneous refinement, after which the best-resolved class was selected for further heterogeneous, homogeneous, non-uniform, and local refinements. The final reconstruction was generated from 227,229 particles with a global resolution of 3.56 Å based on the GSFSC. **B.** Angular distribution of particle orientations used for the cryo-EM reconstruction of ABCG5/G8. The height and color gradient (blue to red) of the cylindrical bars represent the number of particles contributing to each view. **C.** GSFSC curve of the final refined cryo-EM map. **D.** Local resolution map calculated in CryoSPARC and visualized in Chimera, ranging from 3.0 Å (blue) to 5.0 Å (red). **E.** Q-score of the model, mapped per residue, with colors ranging from red (0) to green (0.49; expected Q-score at 3.5 Å resolution) and blue (0.7825; highest Q-score in the model).

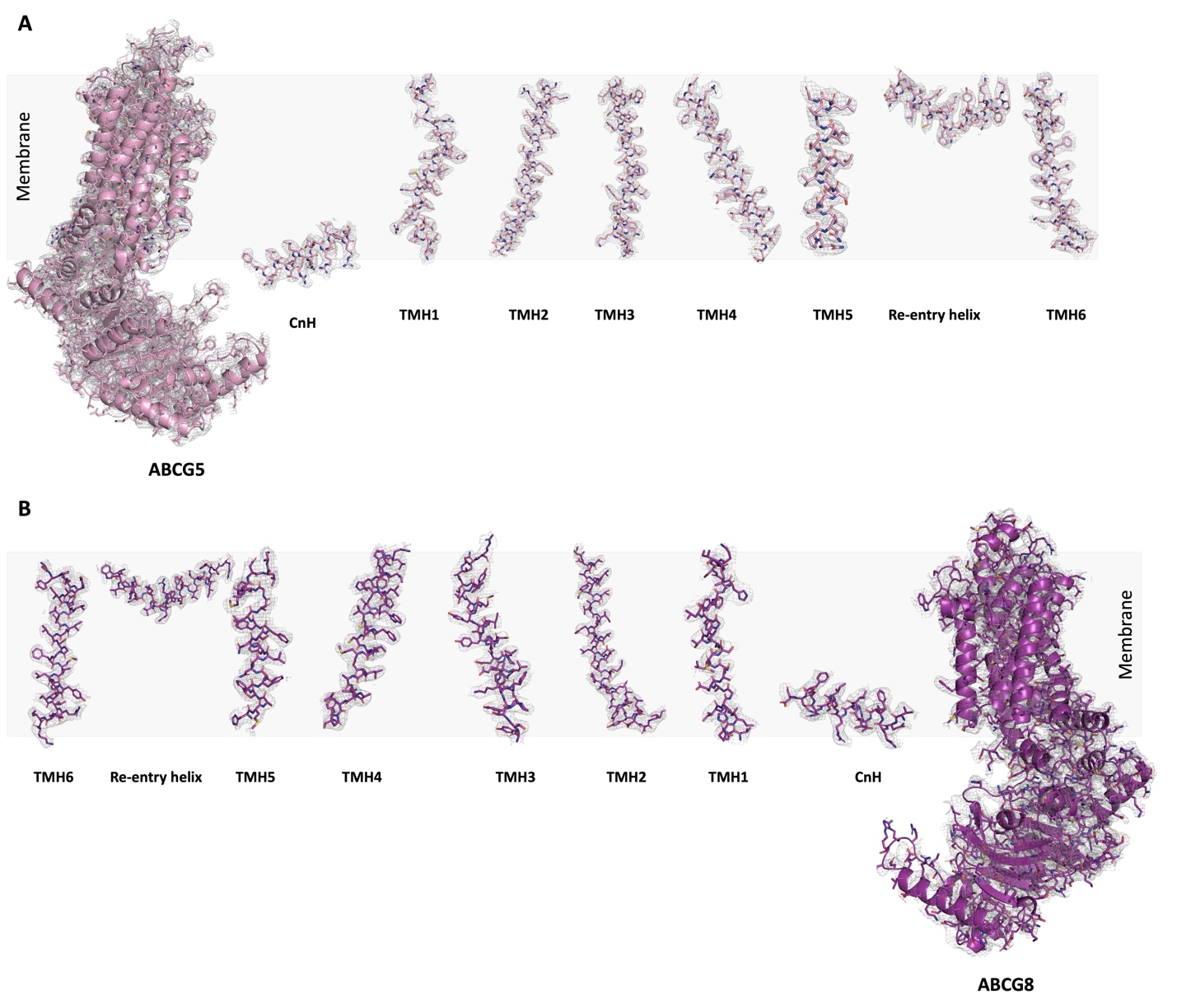

**Figure S3. Refined structural models fitted into the cryo-EM density map.** Cryo-EM density maps with fitted models of ABCG5 (A) and ABCG8 (B), highlighting the TMDs, including TM helices 1–6, the connecting helix (CnH), and the re-entry helix.

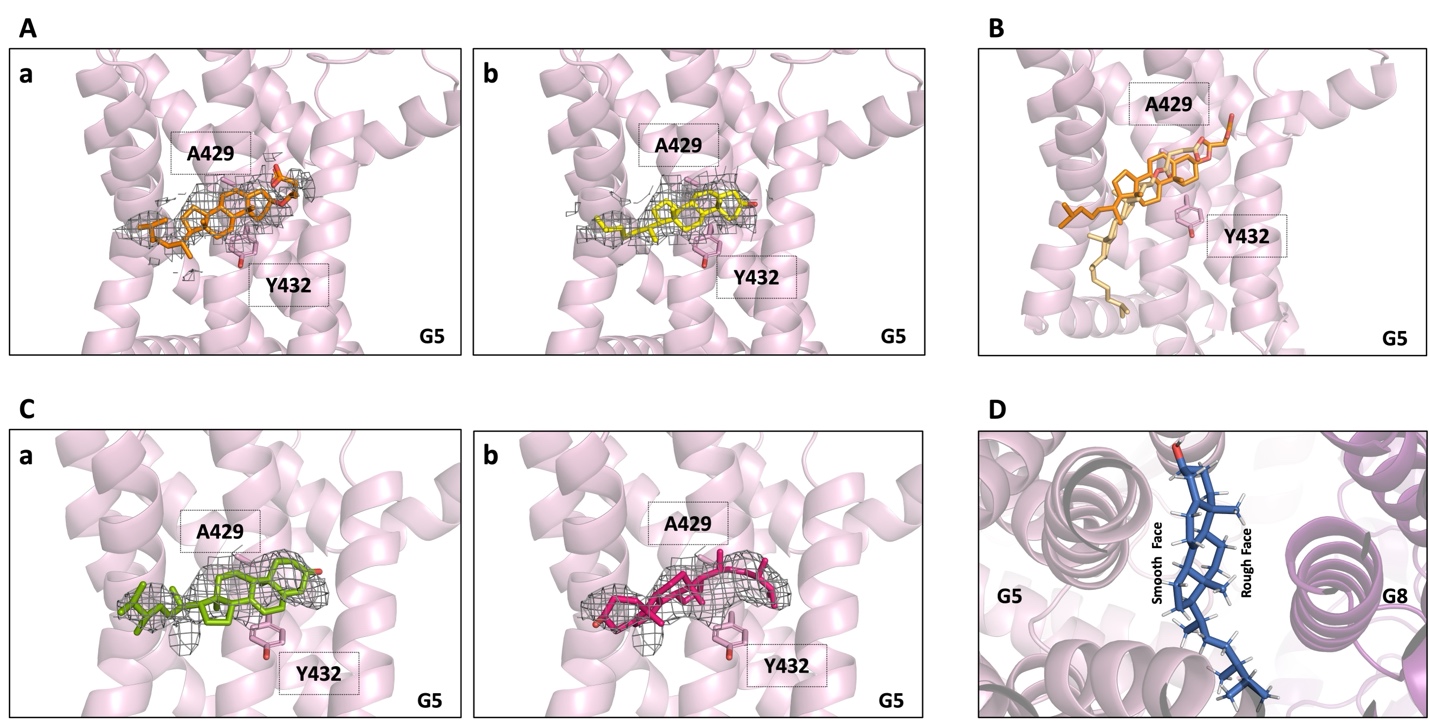

**Figure S4.** **Further analysis of substrate identity and orientation within the observed binding site. A.** CHS (**a**, orange) and cholesterol (**b**, yellow) modeled into the observed density in our cryo-EM map. **B.** Comparison of the initial (orange) and final frames (light orange) from the 200 ns MD simulation of CHS within the ABCG5/G8 binding site. **C.** Proposed orientations of ergosterol within the binding site. **(a)** Ergosterol (green) is positioned such that its β (rough) face interacts with ABCG5, and its α (smooth) face interacts with ABCG8 (CC for ligand in this orientation is 0.63). **(b)** Alternative orientation in which the hydroxyl group of ergosterol (dark pink) is directed toward the surrounding membrane, while the sterane core and hydrocarbon tail extend deeper into the binding cavity (CC for ligand in this orientation is 0.56). **D.** Preferred orientation of ergosterol (blue) within the binding site, with its α (smooth) face interacting with ABCG5 and its β (rough) face interacting with ABCG8. The hydroxyl group is positioned deep within the cavity, while the hydrocarbon tail extends toward the cavity exit (ligand-map CC = 0.8).

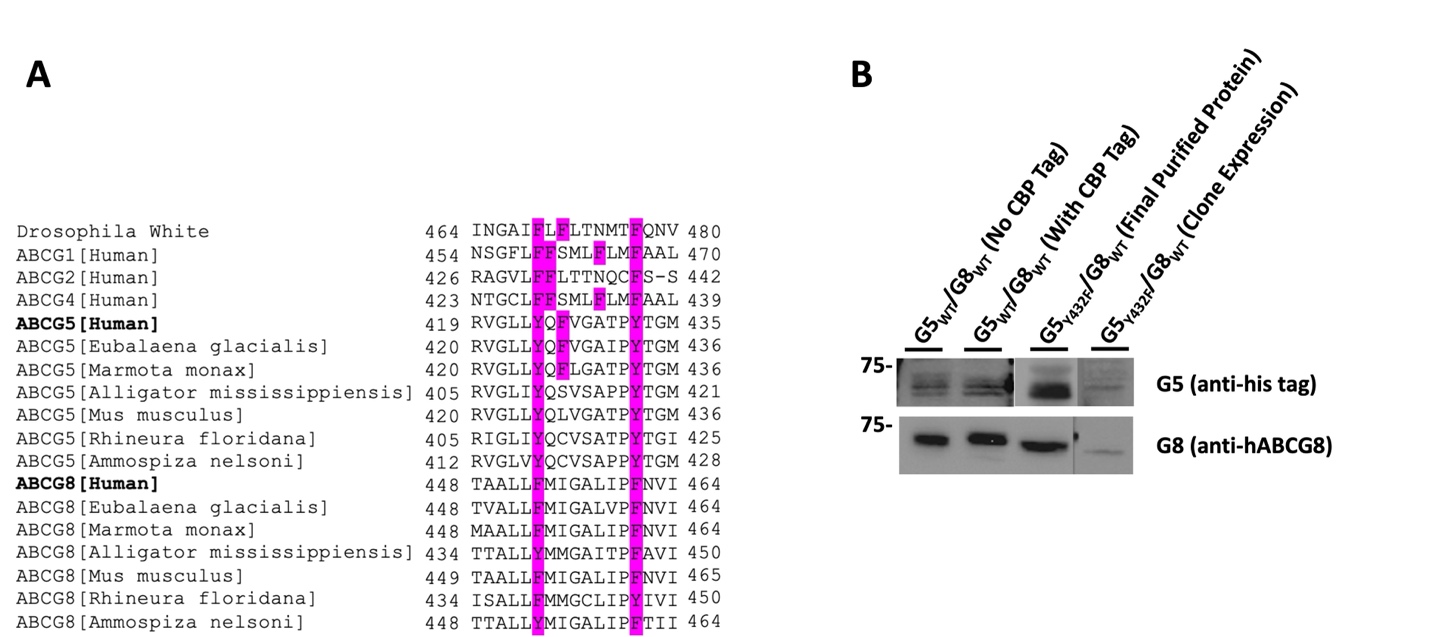

**Figure S5. Sequence conservation of the ABCG aromatic clamp motif and expression analysis of the ABCG5_Y432F_/G8_WT_ mutant. A.** Sequence alignment of human ABCG subfamily members and Drosophila White, as well as ABCG5 and ABCG8 orthologs from six additional species: *Eubalaena glacialis, Marmota monax, Alligator mississippiensis, Mus musculus, Rhineura floridana, and Ammospiza nelsoni*. Residues highlighted in pink correspond to the aromatic clamp motif components conserved across these species and proteins. **B.** Western blot analysis of ABCG5_Y432F_/G8_WT_ mutant and ABCG5_WT_/G8_WT_ constructs. Clonal expression levels and purified protein samples used for ATPase assays were assessed and compared between the mutant and WT constructs.

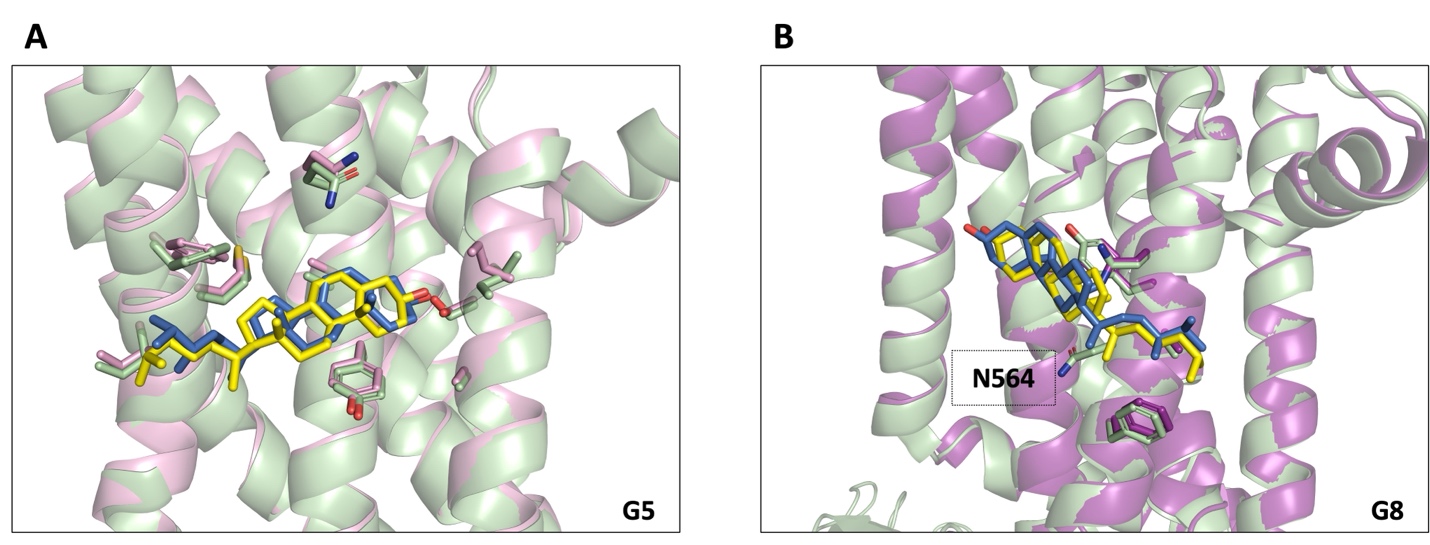

**Figure S6. Structural alignment of ergosterol-bound and cholesterol-bound ABCG5/G8.** The ergosterol-bound cryo-EM structure is shown with ABCG5 in pink, ABCG8 in purple, and ergosterol in blue, whereas the cholesterol-bound structure (PDB ID: 7R8B) is shown in pale green with cholesterol in yellow. **A.** View from the ABCG5 side. **B.** View from the ABCG8 side. ABCG8_N564_ shown in **(B)** contributes to cholesterol binding in the 7R8B structure, but not in the ergosterol binding site.
